# RTEL1 Influences the Abundance and Localization of TERRA RNA

**DOI:** 10.1101/2020.05.12.088583

**Authors:** Fiorella Ghisays, Aitor Garzia, Hexiao Wang, Claudia Canasto-Chibuque, Marcel Hohl, Sharon A. Savage, Thomas Tuschl, John H. J. Petrini

**Affiliations:** Molecular Biology Program, Memorial Sloan-Kettering Cancer Center, New York, New York, United States of America; Laboratory for RNA Molecular Biology, The Rockefeller University, 1230 York Ave, Box 186, New York, NY 10065, USA; Clinical Genetics Branch, Division of Cancer Epidemiology and Genetics, National Cancer Institute, Rockville, MD, USA

## Abstract

Telomere repeat containing RNAs (TERRAs) are a family of long non-coding RNAs transcribed from the sub-telomeric regions of eukaryotic chromosomes. TERRA transcripts can form R-loops at chromosome ends; however the importance of these structures or the regulation of TERRA expression and retention in telomeric R-loops remain unclear. Here, we show that the RTEL1 (Regulator of Telomere Length 1) helicase influences the abundance and localization of TERRA in human cells. Depletion of RTEL1 leads to increased levels of TERRA RNA while reducing TERRA-containing R loops at telomeres. *In vitro*, RTEL1 shows a strong preference for binding G-quadruplex structures which form in TERRA. This binding is mediated by the C-terminal region of RTEL1, and is independent of the RTEL1 helicase domain. RTEL1 binding to TERRA appears to be essential for cell viability, underscoring the importance of this function. Degradation of TERRA containing R-loops by overexpression of RNAse H1 partially recapitulates the increased TERRA levels and telomeric instability associated with RTEL1 deficiency. Collectively, these data suggest that regulation of TERRA at the telomeres is a key function of the RTEL1 helicase, and that loss of that function may contribute to the disease phenotypes of patients with *RTEL1* mutations.

## Introduction

RTEL1 is an (Fe-S) cluster containing helicase that belongs to the DEAH superfamily 2 (SF2) and specifies 5’-3’ helicase activity (Uringa et al., 2011; Wu and Brosh, 2012). The *RTEL1* gene was first identified in a four-gene cluster on chromosome 20 found to be amplified in gastrointestinal cancers (then called NHL) (Bai et al., 2000). The gene was subsequently identified as a regulator of telomere length in the mouse and renamed RTEL1 (for **R**egulator of **Te**lomere **L**ength) (Ding et al., 2004).

RTEL1 plays diverse functional roles that bear upon several aspects of genome integrity and gene expression. RTEL1 influences telomere stability by facilitating replication at telomeres, likely by removing secondary structures that would otherwise inhibit replication fork progression (Margalef et al., 2018; Sarek et al., 2015). In addition, it influences genome stability via its roles in regulating DNA replication and DNA recombination (Vannier et al., 2013). More recently, RTEL1 has been shown to disrupt R-loops (RNA:DNA hybrid structures) in GC-rich regions of the genome, and to promote MiDAS, a mitotic DNA synthesis process that occurs in common fragile sites and other late replicating genomic loci (Minocherhomji et al., 2015; Wu, 2020). Finally, the protein also appears to influence RNA trafficking, as cells from patients with inherited *RTEL1* mutations exhibit aberrant localization of pre-U2 RNA and associated ribonucleoproteins (RNPs) (Schertzer et al., 2015).

Germline mutations in *RTEL1* underlie Dyskeratosis Congenita (DC) and the more severe variant, Hoyeraal-Hreidarsson syndrome (HH) (Bertuch, 2016; Niewisch and Savage, 2019). At the cellular level, these diseases are associated with very short and heterogeneous telomeres, genomic instability, and sensitivity to clastogens, consistent with the role of RTEL1 in preserving genome integrity (Ballew et al., 2013a; Faure et al., 2013). Some disease-causing *RTEL1* alleles exhibit autosomal dominant behavior, suggesting that the protein may function in a dimeric or multimeric assembly (Ballew et al., 2013b; Borie et al., 2019; Newton et al., 2016; Touzot et al., 2016).

Telomeric repeat containing RNAs (TERRAs) are RNA polymerase II transcripts emanating from subtelomeric regions that extend into the TTAGGG telomeric repeats (Azzalin et al., 2007; Fedick et al., 2015; Feretzaki et al., 2019; Schoeftner and Blasco, 2008). TERRA transcripts are heterogeneous in size and can be associated with chromosome ends by forming R-loops either co-transcriptionally or post-transcriptionally. In the latter circumstance, the TERRA RNA may be recruited via binding to telomeric proteins (Aguilera and Garcia-Muse, 2012; Azzalin et al., 2007; Balk et al., 2013; Chu et al., 2017; Feretzaki et al., 2019; Lee et al., 2018; Lopez de Silanes et al., 2010; Porro et al., 2014).

TERRA is transcribed from at least a subset of chromosome ends, and the mechanisms of regulation may differ between chromosomes. TERRA binds to telomeres as well as interstitial sites where it can influence gene expression (Beishline et al., 2017; Chu et al., 2017; Feretzaki et al., 2019; Sagie et al., 2017). Despite this broad distribution of binding sites, the most prominent role of TERRA is at the telomere. Depletion of TERRA leads to myriad indices of telomere dysfunction such as the loss of telomeric DNA, the formation of telomere damage associated foci (TIFs) (Takai et al., 2003), and telomere driven chromosome aberrations. Notably, TERRA interacts with, and appears to antagonize the functions of the chromatin remodeler ATRX, which plays a key role in modulating telomeric chromatin structure (Chu et al., 2017; Eid et al., 2015; Nguyen et al., 2017).

Due to its G-rich sequence, TERRA can form stable, four-stranded structures known as G-quadruplexes *in vitro* (Bao et al., 2017; Biffi et al., 2012; Sen and Gilbert, 1988). G-quadruplex structures can form in both G-rich DNA and RNA sequences *in vitro* under physiological conditions, and are likely to constitute barriers to protein translocation during DNA replication, transcription, and mRNA translation if present in cells (Hansel-Hertsch et al., 2017). There are dedicated protein machineries with the capacity to unfold G-quadruplex structures in mammalian cells (Guo and Bartel, 2016). However, it is not clear whether TERRA forms G-quadruplex structures *in vivo* and if so, whether those structures are biologically relevant.

Here we present evidence that RTEL1 influences the levels and localization of TERRA via direct physical interaction. We identified an RNA binding domain in the RTEL1 C-terminus that exhibits a strong preference for G-quadruplex folded TERRA RNA over unfolded RNA. RTEL1 deficiency causes a dramatic increase in TERRA levels, but reduced localization of TERRA RNA to chromosome ends. Our data thus reveal a previously unrecognized function of RTEL1 in regulating the disposition of the TERRA RNA. The data are consistent with the observation that misregulation of TERRA compromises telomere maintenance, and suggest a novel mechanism by which *RTEL1* hypomorphism may cause telomere instability and contribute to the clinical phenotypes of DC and HH (Arora et al., 2014; Beishline et al., 2017; Chu et al., 2017; Cusanelli et al., 2013; Deng et al., 2012; Graf et al., 2017; Lopez de Silanes et al., 2014; Moravec et al., 2016; Pfeiffer and Lingner, 2012; Porro et al., 2014).

## Results

Homozygosity of the *RTEL1*^*R1264H*^ pathogenic variant causes HH, which occurs as a heterozygous founder mutation in the Ashkenazi Jewish population (Ballew et al., 2013a). The mutation falls within the RING domain at the protein’s C-terminus, and its implication as an underlying cause of HH provided the first evidence of the RING domain’s functional significance (Ballew et al., 2013a). Typically, the RING domain structure is dependent upon the coordination of Zn^2+^ atoms via cysteine and histidine residues. The R→H change in the *RTEL1*^*R1264H*^ RING domain introduces an additional potential Zn^2+^ coordination partner that may alter the RING domain’s structure.

### Domain Analysis of RTEL1

To examine the functional consequences of the *RTEL1*^*R1264H*^ mutation at the molecular level, we designed a series of constructs for the production of various RTEL1 protein domains (**Figure 1A**). First, differentially tagged (myc and FLAG) wild type and *RTEL1*^*R1264H*^ full length cDNAs were co-expressed in HEK293 cells. FLAG-tagged RTEL1 was recovered in myc immunoprecipitates of both *RTEL1*^*R1264H*^ and wild type RTEL1, suggesting that the protein can form a higher order assembly (**Figure 1B, left**).

**Figure 1.**
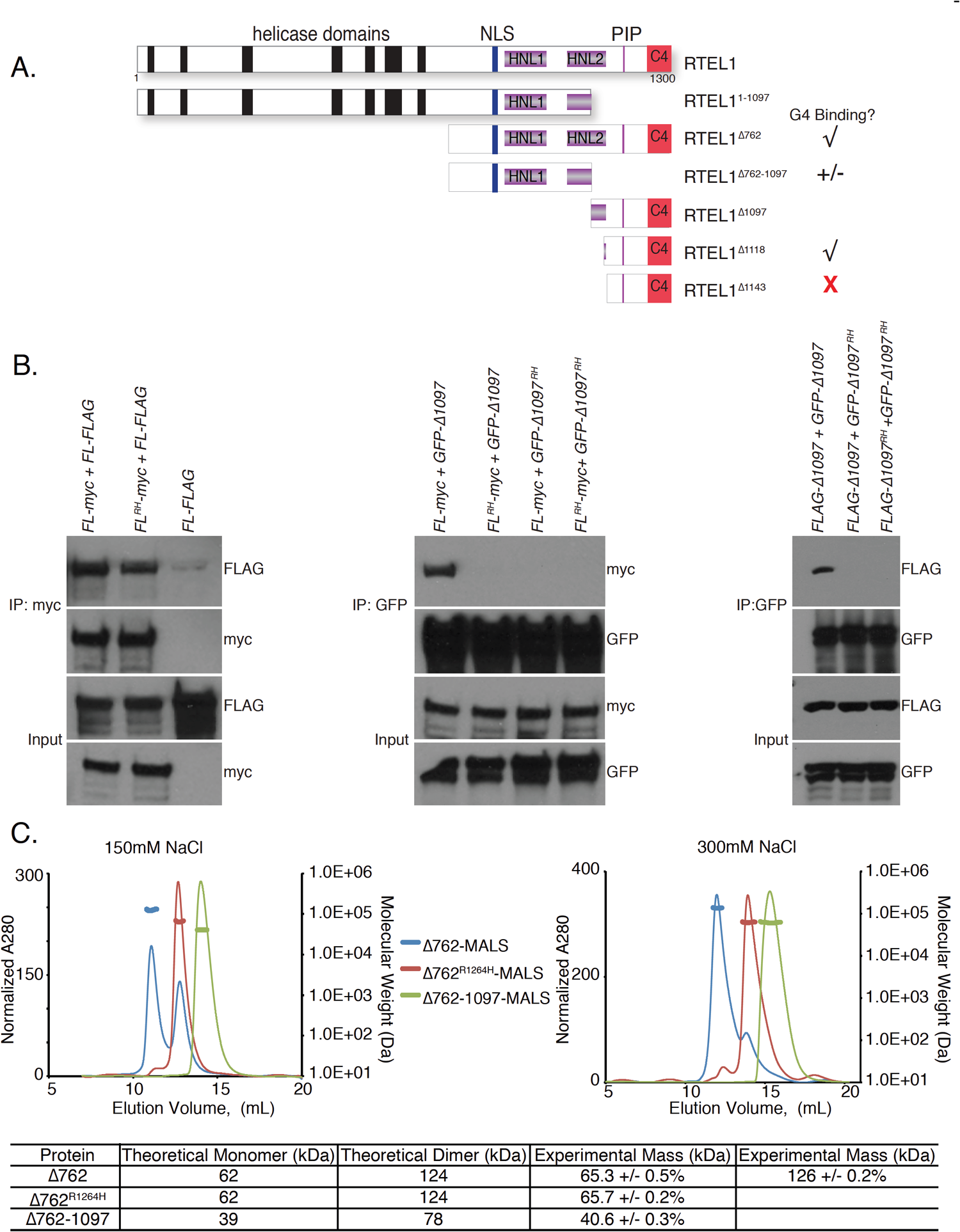
Domain Analysis of RTEL1. (A) Schematic representation of RTEL1 proteins and constructs used in this study with G4 binding ability illustrated where applicable. (B) RTEL1 interacts with itself in cells and the *RTEL1*^*R1264H*^ mutation disrupts interactions within the RING domain. Left, Myc immunoprecipitations from HEK293T cells co-expressing FLAG and myc-tagged *RTEL1 (FL)* and *RTEL1*^*R1264H*^ (*FL* ^*RH*^) were carried out. Immunoblotting with FLAG indicates co-IP of both *RTEL1* and *RTEL1*^*R1264H*^. Center, immunoblotting with myc indicates co-IP of RTEL1^Δ1097^ (Δ1097), but not RTEL1^Δ1097^ ^R1264H^ (Δ1097 ^RH^) with GFP-tagged *RTEL1* and *RTEL1*^*R1264H*^. Right, immunoblotting with FLAG shows co-IP of RTEL1^Δ1097^, but not RTEL1^Δ1097R1264H^. (C) Oligomeric states of RTEL1 proteins were determined by SEC-MALS analysis of RTEL1^Δ762^ (blue), RTEL1^Δ762R1264H^ (red), and RTEL1^Δ762-1097^ (green) at 150 and 300 mM NaCl. Normalized A280 are shown and the calculated molecular mass is shown as a line in the corresponding color across each peak with the secondary scale on the right. The expected and calculated molecular weights are shown. (See also Figure S1).

This assembly is mediated by spatially distinct regions of RTEL1, one in the N-terminal helicase domain and the other in the C-terminal RING domain. The protein product of *FLAG*-*RTEL1*^*Δ1097*^, which comprises the C-terminal 203 amino acids of RTEL1 including the PIP box and RING domain (**Figure 1A**), co-immunoprecipitated with both full length RTEL1 and itself. These interactions were abolished by the *RTEL1*^*R1264H*^ mutation (**Figure 1B, center and right**), indicating that the RING domain constitutes an independent homotypic interaction domain that is rendered non-functional by the *RTEL1*^*R1264H*^ mutation. Full length RTEL1^R1264H^ retains the ability to interact with wild type full length RTEL1, indicating that the N-terminal half of RTEL1 also promotes oligomerization (**Figure 1B, left**).

To characterize the C-terminal oligomerization domain of RTEL1, we began with RTEL1^Δ762^ (which lacks the helicase domain but retains the harmonin N-like, PIP box, and RING domains of RTEL1), and designed variants, including the *RTEL1*^Δ762*R1264H*^ mutation and various deletions within RTEL1^Δ762^ that removed the RING domain and additional proximal sequences (**Figure 1A**). We purified recombinant His-tagged RTEL1^Δ762^ from *E*.*coli*. Size exclusion chromatography (SEC) revealed that RTEL1^Δ762^ was present in two monodispersed peaks (**Figure 1C**). Coupling the SEC with multiple angle light scattering (SEC-MALS) showed the molecular masses of the RTEL1^Δ762^ peaks to be 65.3 ± 0.5% and 126 ± 0.2% kDa respectively corresponding to the expected monomer and dimer masses of RTEL1^Δ762^ (62kDa and 124kDa) (**Figure 1C**). The dimer peak was virtually absent in the RTEL1^Δ762R1264H^ protein and the calculated molecular mass was 65.7 ± 0.2% kDa, corresponding to that of a monomer (**Figure 1C**). The RTEL1^Δ762-1097^ protein, which lacks the RING domain, is also a monomer (**Figure 1C**).

SEC-MALS experiments showed that the RTEL1^Δ762^ dimeric species was favored at higher ionic strength. Elevation of the salt concentration from 150 mM to 300 mM NaCl favored the RTEL1^Δ762^ dimeric species but had no effect on the oligomeric state of either RTEL1^Δ762R1264H^ or RTEL1^Δ762-1097^ proteins (**Figure 1C**). These results suggest that dimerization of the C-terminal RTEL1 domain is dependent on the RING domain in combination with hydrophobic interactions that are stabilized at higher ionic strength. Hence, the RING domain of RTEL1 is a dimerization domain and the *RTEL1*^*R1264H*^ mutation disrupts that function *in vitro* and in cells.

### Heterotypic Interactions at the RTEL1 C-terminus

Having identified the homotypic interaction in the C-terminus of RTEL1, we reasoned that heterotypic protein interactions at the RTEL1 C-terminus could also provide insight regarding helicase independent functions of RTEL1, and shed light on the function(s) of the RING domain as well as the consequences of the *RTEL1*^*R1264H*^ mutation. We used a BioID proximity dependent labeling system (Roux et al., 2012) to identify RTEL1 C-terminal interacting proteins. This system uses the promiscuous biotin ligase BirA^R118G^ fused to the protein of interest. Upon addition of biotin to cells expressing BirA fusion proteins, biotin is covalently linked via lysine residues within a 10 nm radius of the BirA^R118G^ tag (Kim et al., 2014), and biotinylated proteins can subsequently be purified via streptavidin binding. The biotin-labeled proteins can then be identified by mass spectrometry (Roux et al., 2012).

BirA^R118G^ was fused to the C-terminus of full length RTEL1 and RTEL1^1-1097^, from which part of the second Harmonin N-like repeat, the PIP box, and the RING domain are deleted (**See Figure 1A and S1A**). RTEL1 is normally located in the nucleus and fusion to BirA did not change its intracellular localization (**Figure S1B**). Nevertheless, the biotin-labeled proteins identified in this experiment included numerous RNA binding proteins that function in the cytoplasm. These RNA binding proteins were enriched by interaction with full-length RTEL1 relative to RTEL1^1-1097^. Eighteen proteins obtained in our screen were also recovered in a screen that identified 38 proteins that bound the telomeric RNA, TERRA (Lopez de Silanes et al., 2010) (**Figure S1C**). Additionally, SFPQ and NONO have been recently characterized as TERRA binding proteins and were among the top hits in our screen (Petti et al., 2019). RTEL1 was also found in a separate screen for TERRA interacting proteins (Chu et al., 2017).

### The RTEL1 C-terminal Domain Binds TERRA and Other G-quadruplex Structures

We did not observe direct interactions between RTEL1 and the TERRA binding proteins previously identified (data not shown). Instead, we found that RTEL1 bound TERRA directly *in vitro* via a domain in the C-terminus. We assayed binding of RTEL1^Δ762^ (**Figure 1A**) to TERRA by fluorescence anisotropy. First, we determined the dissociation constant (K_D_) for fluorescein labeled TERRA RNAs that fold into G-quadruplex structures (TERRA), a mutant TERRA RNA that is unable to form the G-quadruplex structure (TERRA-MUT), and TERRA RNA that had not been folded into a G-quadruplex (**Figure 2A**). TERRA RNA and the analogous DNA molecules were folded into G-quadruplexes by heating the oligonucleotides to 95°C then slowly cooling them in 50 mM KCl, and validated by circular dichroism as previously described (Biffi et al., 2012; Yangyuoru et al., 2013). Two RNAs unrelated to TERRA that were either unstructured (AU rich) or differently structured (PolyA) were also examined (Safaee et al., 2013) (**Figure 2B**). 50 nM RNA was incubated with increasing concentrations of RTEL1^Δ762^ (0-5 *µ*M) and incubated for 30 minutes at RT. RTEL1^Δ762^ bound to all tested RNAs and showed a striking preference for the folded TERRA RNA, exhibiting a 500-fold preference over the PolyA RNA, and 100-fold over the TERRA-MUT substrate incapable of forming G-quadruplexes (**Figure 2B and S2A**).

**Figure 2.**
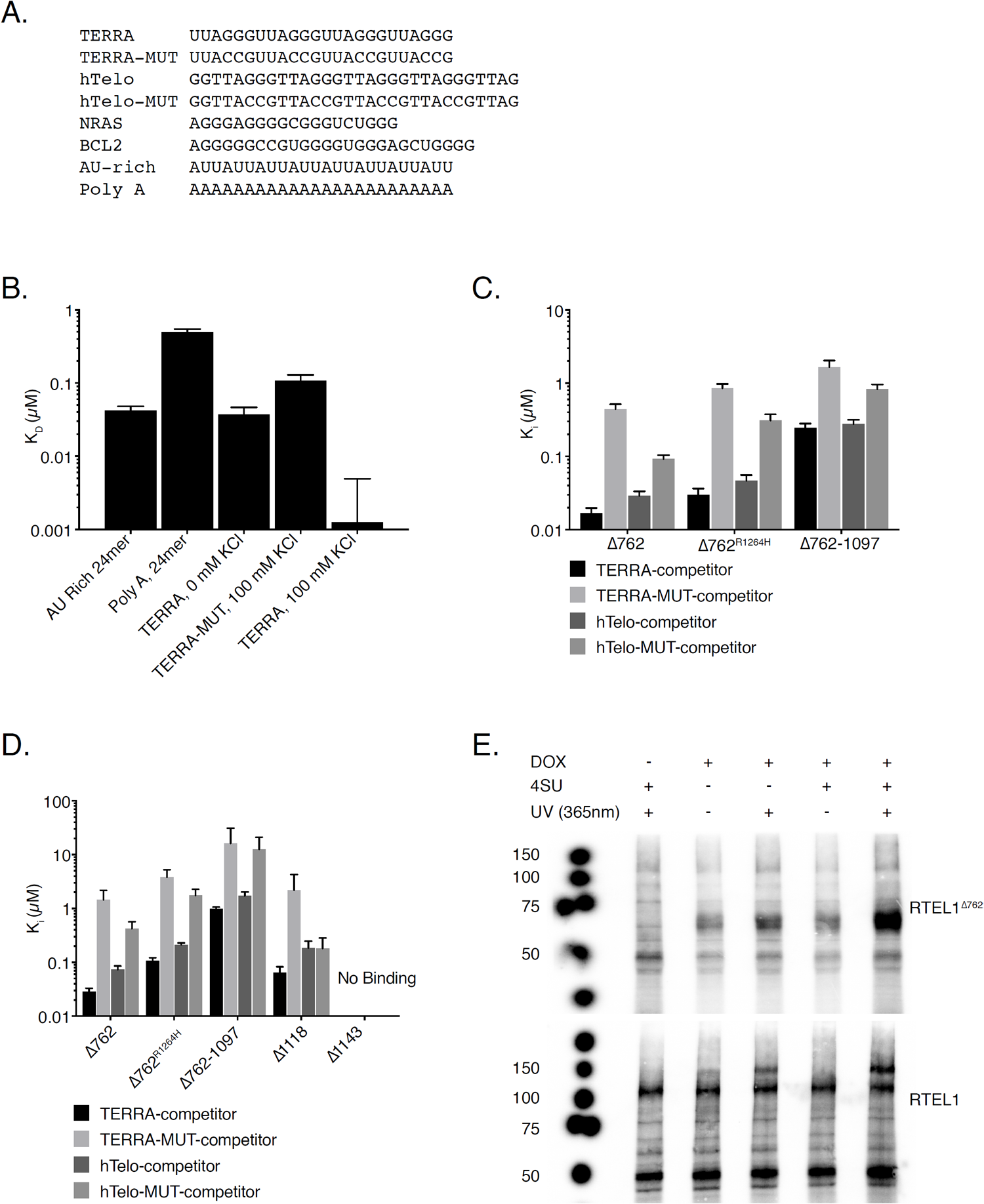
The RTEL1 C-terminal Domain Binds TERRA and Other G-quadruplex Structures. (A) Sequences of DNA and RNA substrates used in this study. (B) Nucleic acid binding of RTEL1 proteins lacking the helicase domains was monitored by fluorescence anisotropy. Bar graph shows dissociation constants (Kd) for TERRA and TERRA-MUT RNAs folded in the presence and absence of KCl and, AU-rich and polyA RNA controls. (C) Increasing concentrations of the indicated oligonucleotides were added to reactions containing RTEL1 proteins and a 24mer FAM-TERRA-MUT RNA at 200 nM. Bar graphs depict apparent dissociation constants (Ki) derived by competition of the bound FAM-TERRA-MUT by the indicated oligonucleotides using fluorescence anisotropy. (D) Ki’s of the indicated RTEL1 deletion constructs were derived by competition studies of FAM-TERRA-MUT and indicated RTEL1 proteins at 1 *µ*M as in panel C. No binding signal was observed for the RTEL1^Δ1143^ protein. All binding assays were conducted in triplicate, mean and standard deviation are shown. (E) Autoradiograph of cross-linked, ^32^P-labelled RNA-FLAG-tagged-RTEL1 immunoprecipitates. FLAG-tagged RTEL1 and RTEL1^Δ762^ proteins were separated by SDS-PAGE after 4SU PAR-CLIP. (See also Figure S2-S6).

The K_D_ for RTEL1^Δ762^ binding to TERRA (< 10 nM) is at the detection limit for anisotropy, which may compromise the accuracy of the measurement. As an alternative to direct determination of K_D_, we again used fluorescence anisotropy to measure the ability of substrates to compete with TERRA-MUT RNA and thereby derive Ki values as a surrogate for the apparent K_D_. RTEL1^Δ762^, RTEL1^Δ762R1264H,^ and RTEL1^Δ762-1097^ proteins (**Figure 1A**) were pre-incubated with TERRA-MUT RNA (**Figure 2A**) for 30 minutes to allow protein-RNA complexes to form, followed by addition of increasing amounts of wild type TERRA RNA or DNA competitor substrates. The K_i_ of RTEL1^Δ762^ for the folded TERRA RNA was 17± 3 nM, 26-fold greater affinity than binding to TERRA-MUT RNA (K_i_ 442 ±. 74) (**Figure 2C and S2B**).

RTEL1 bound G strand telomeric DNA (hTelo) with similar affinity to TERRA RNA; however, the preference for folded telomeric DNA relative to unfolded DNA (hTelo-MUT) is diminished by approximately three fold. The K_i_ for RTEL1^Δ762^ binding hTelo DNA was 29 ± 4 nM, and for hTelo-MUT DNA (K_i_ 93 ± 11 nM) (**Figure 2C and S2B**). Hence, this domain of RTEL1 discriminates G4 from other RNA structures to greater extent than the same structures in DNA. Within the context of telomeric chromatin, it is unclear whether G4 DNA and RNA structures co-exist, but the selectivity of RTEL1 for folded RNA may be relevant to its telomeric functions.

### The G4 Binding Domain of RTEL1

To identify the location of the G4 binding domain in the RTEL1 C-terminus, we used fluorescence anisotropy as above with fragments of RTEL1^Δ762^ (**Figure1A**). The RTEL1^Δ762R1264H^ protein displayed similar binding affinities to the wild type RTEL1^Δ762^ protein with only a two to four-fold difference in Ki values (**Figure 2C and S3**). These data indicate that RNA/DNA binding is not directly mediated by the RING domain. The decreased apparent affinity of RTEL1^Δ762R1264H^ likely reflects the difference in valency of the TERRA binding domain in monomeric *vs*. dimeric RTEL1^Δ762^.

Deletion of the C-terminal 230 residues of Δ762 (resulting in RTEL1^Δ762-1097^ protein) reduced binding affinities by approximately 15-fold (**Figure 2C and S2B**) and led to a reduced preference for TERRA—6.7 fold relative to the 26.4-fold preference for TERRA in the wild type protein. Unlike binding to RTEL1^Δ762^, the residual binding activity of RTEL1^Δ762-1097^ was acutely sensitive to salt concentration (**Figure S4**). The binding affinity and selectivity for G4 binding of the RTEL1^Δ1118^ protein product were similar to RTEL1^Δ762^ (**Figure 2D**), whereas binding was largely absent for RTEL1^Δ1143^ (**Figure 2D, S3 and S5**). These data suggest that G4 binding is primarily specified by residues between amino acid 1097 and the RING domain (**Figure 1A**).

As RTEL1 functions affect interstitial as well as telomeric chromosomal sites, we asked whether RTEL1 bound to G4 structures forming in RNA encoded from interstitial loci. We used two RNAs predicted to form G4 structures that fall within the 5’ untranslated regions of NRAS and BCL2 (**Figure 2A**) (Biffi et al., 2012). RTEL1^Δ762^, RTEL1^Δ762R1264H,^ and RTEL1^Δ762-1097^ proteins bound both NRAS and BCL2 RNAs with similar affinities to TERRA (**Figure S6A and S6B**).

### RTEL1 Binds RNA In Cells

To determine whether RTEL1 bound RNA in cells, we performed performed 4-thiouridine (4SU) photoactivatable ribonucleoside-enhanced cross-linking and immunoprecipitation (PAR-CLIP) (Garzia et al., 2017b). FLAG immunoprecipitation from HEK293 cells overexpressing FLAG-HA-tagged RTEL1 and RTEL1^Δ762^ was performed followed by RNAse treatment. RNA within immunoprecipitated complexes was 5’ end-radiolabeled with polynucleotide kinase and visualized by autoradiography. Both full length RTEL1 and RTEL1^Δ762^ immunoprecipitates contained radiolabeled RNA (**Figure 2E and S6C**). We were unable to obtain reliable RNA sequencing data from the material in the RTEL1 PAR-CLIP, possibly due to the structure of the RNA(s) bound in that context.

### RTEL1 Influences TERRA Levels

To understand the functional significance of RTEL1 binding to TERRA, we set out to assess the disposition of TERRA in the context of RTEL1 deficiency. *RTEL1* was inactivated via CRISPR-Cas9-mediated deletion of exon #2 in HEK293 cells (hereafter, RTEL1-KO) (Ran et al., 2013) (**Figure 3A and 3B**). The effect of RTEL1 deficiency on telomere stability was examined by fluorescence in situ hybridization (FISH) using a PNA-[TTAAGGG]^3^ probe (Tel FISH). RTEL1-KO cells displayed a significant loss of telomere signal from one or both sister chromatids when compared to wild type HEK293 cells (48.9% ±11.6 *vs*. 19.4% ± 6.2) (**Figure 3C**), consistent with previous results in both human and mouse cells (Ballew et al., 2013a; Le Guen et al., 2013; Sarek et al., 2015; Uringa et al., 2011; Vannier et al., 2012).

**Figure 3.**
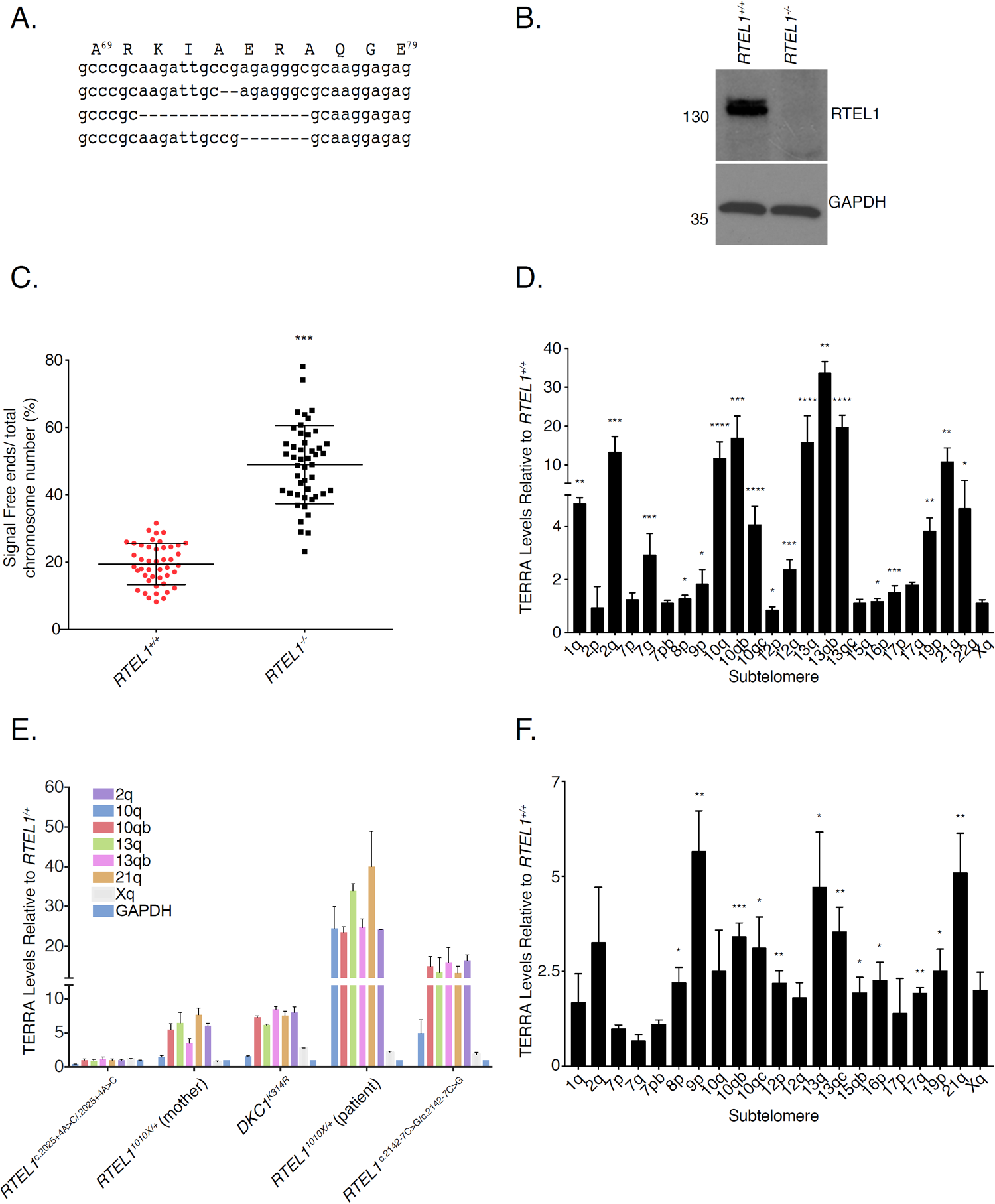
RTEL1 Influences TERRA Levels. (A) Sequence alignment of the PCR products amplified from the HEK293 cells genomic segment of *RTEL1* targeted by sgRNA revealing disruption of the coding sequence (exon #2 of *RTEL1)*. (B) Western blot of RTEL1 protein levels in wild type and *RTEL1* knockout HEK293 cells. (C) Quantification of telomere loss per chromosome. Data represents the average of at least 50 metaphases as mean and standard deviation (***p < 0.001, t-test). (D) TERRA levels at specific chromosome ends are elevated in RTEL1 deficient cells. Bars represent the relative TERRA expression measured by qRT-PCR at eighteen chromosome ends. Values are sample averages of at least 4 experimental repeats (*p < 0.05, **p <0.01, ***p < 0.001, t-test). (E) TERRA levels are elevated in cells from a panel of patient derived LCL lines. TERRA levels were measured by qRT-PCR. Values are sample averages of at least 3 experimental repeats (*p < 0.05, **p <0.01, ***p < 0.001, t-test). (F) TERRA levels are elevated in a fibroblast cell line derived from a HHS patient homozygous for the *RTEL1*^*R1264H*^ allele. TERRA levels were measured by qRT-PCR. Values are sample averages of at least 3 experimental repeats (*p < 0.05, **p <0.01, ***p < 0.001, t-test). (See also Figure S6 and S7).

We next examined the levels of TERRA RNA. TERRA transcription initiates at subtelomeric regions and extends into the telomeric repeats. The levels of TERRA transcripts originating at various chromosome ends was measured by quantitative reverse transcription PCR (qRT-PCR) using primers derived from the respective subtelomeric sequences (Deng et al., 2012; Feretzaki and Lingner, 2017; Sagie et al., 2017) (**Table S1**). We tested 18 chromosome ends, some with more than one primer set. We found that TERRA levels were markedly elevated in RTEL1-KO cells when compared to wild type cells. The fold change was highest for chromosome ends 2q, 10q, 13q, and 21q (**Figure 3D**). These results are consistent with the view that TERRA transcription is controlled by different factors at different chromosome ends (Deng et al., 2012; Feretzaki et al., 2019). RTEL1 deficiency did not affect the levels of mRNA transcribed from genes that are known to form R-loops or that of *NRAS* and *BCL2* mRNAs, the 5’ UTRs of which form G-quadruplexes that are bound by RTEL1 *in vitro* (**Figure S6D**).

TERRA levels were also elevated in a panel of lymphoblastoid cell lines derived from HH and DC patients as well as a fibroblast cell line homozygous for the *RTEL1*^*R1264H*^ mutation. The lymphoblastoid lines examined were derived from a patient with HH harboring an autosomal dominant mutation, *RTEL1*^*1010x/+*^ (NCI-164-1), the patient’s mother who presented with short telomeres but it is clinically unaffected (NCI-164-3), two cell lines from patients with HH due to with homozygous *RTEL1* splice variants (*RTEL1 c*.*2025+4A>C* (NCI-323-1) and *RTEL1 c*.*2142-7C>G* (NCI-347-1)), and a DC patient cell line with a dyskerin mutation (*DKC1*^*K314R*^ (NCI-106-3)) (Ballew et al., 2013a; Ballew et al., 2013b; Ungar et al., 2018). Cells derived from a healthy donor proven negative for telomere biology gene mutations by exome sequencing (NCI-165-2) were used as the wild type control for lymphoblastoid cell lines, and a laboratory stock of previously immortalized BJ-hTERT’s were used as controls for *RTEL1*^*R1264H*^ fibroblast cell lines as no healthy donor samples were available.

TERRA transcripts emanating from the same subtelomeric regions as in the RTEL1-KO line (*i*.*e*., 2q, 10q, 13q, and 21q) were elevated in cells derived from the *RTEL1*^*1010x/+*^ proband, the proband’s mother, the *RTEL1 c*.*2142-7C>G*, and the *DKC1*^*K314R*^ patient samples (**Figure 3E and S7C-S7E**). TERRA levels were also significantly elevated in a fibroblast cell line derived from a HH patient homozygous for the *RTEL1*^*R1264H*^ allele (Ballew et al., 2013a) (**Figure 3F**). Conversely, TERRA levels were normal in the cells with the *RTEL1 c*.*2025+4A>C* mutation in which RTEL1 protein levels were not affected (**Figure S7B**), and TERRA transcripts emanating from chromosome 17p, which are regulated by CTCF binding (Beishline et al., 2017) were elevated in the *DKC1*^*K314R*^ and *RTEL1 c*.*2142-7C>G* cell lines (**Figure S7D**). 2q, 10q, 13q, and 21q transcripts were unaffected in that cell line. These data suggest that alterations in TERRA levels associated with defects in RTEL1 may contribute to the disease phenotypes in HH and in DC.

In our hands, the RTEL1-KO cell line exhibits limited growth capacity and becomes senescent within one to two months, consistent with previous data (Barber et al., 2008). Complementation with *RTEL1*^*1-762*^, *RTEL1*^*1-1118*^, *RTEL1*^*Δ762*^, and *RTEL1*^*R1264H*^ (**Figure 1A**), failed to rescue the senescence phenotype, suggesting that both the TERRA binding domain and the helicase domain are required for viability. However, clones were established upon complementation with wild type *RTEL1* and the helicase deficient *RTEL1*^*K48R*^ mutant (**Figure S7A and data not shown**), suggesting that helicase activity *per se* is not essential.

Complementation of RTEL1-KO cells normalized TERRA levels in three independent transductions. Following introduction of wild type *FLAG-RTEL1*, TERRA levels decreased approximately ten-fold from all chromosomes tested (**Figure 4A and Figure S7A**). TERRA levels were also reduced in *FLAG-RTEL1*^*K48R*^-expressing clones but remained significantly higher than cells complemented with wild type *RTEL1* (**Figure 4A**). RNA-seq analyses of wild type, RTEL1-KO HEK293 cells and cells complemented with *FLAG-RTEL1* or *FLAG-RTEL1*^*K48R*^ showed that subtelomeric read counts in the RTEL1-KO were elevated and reduced upon complementation with *FLAG-RTEL1 or FLAG-RTEL1*^*K48R*^ (**Figure 4B**). The magnitude of the increased read counts is less than observed in Q-PCR analyses, likely reflecting that TERRA increases were greatest in four chromosome ends; RNA-seq essentially averages high and low expression levels of TERRA coming from different chromosome ends. These results suggest that RTEL1 influences TERRA levels in a manner that is partially helicase dependent.

**Figure 4.**
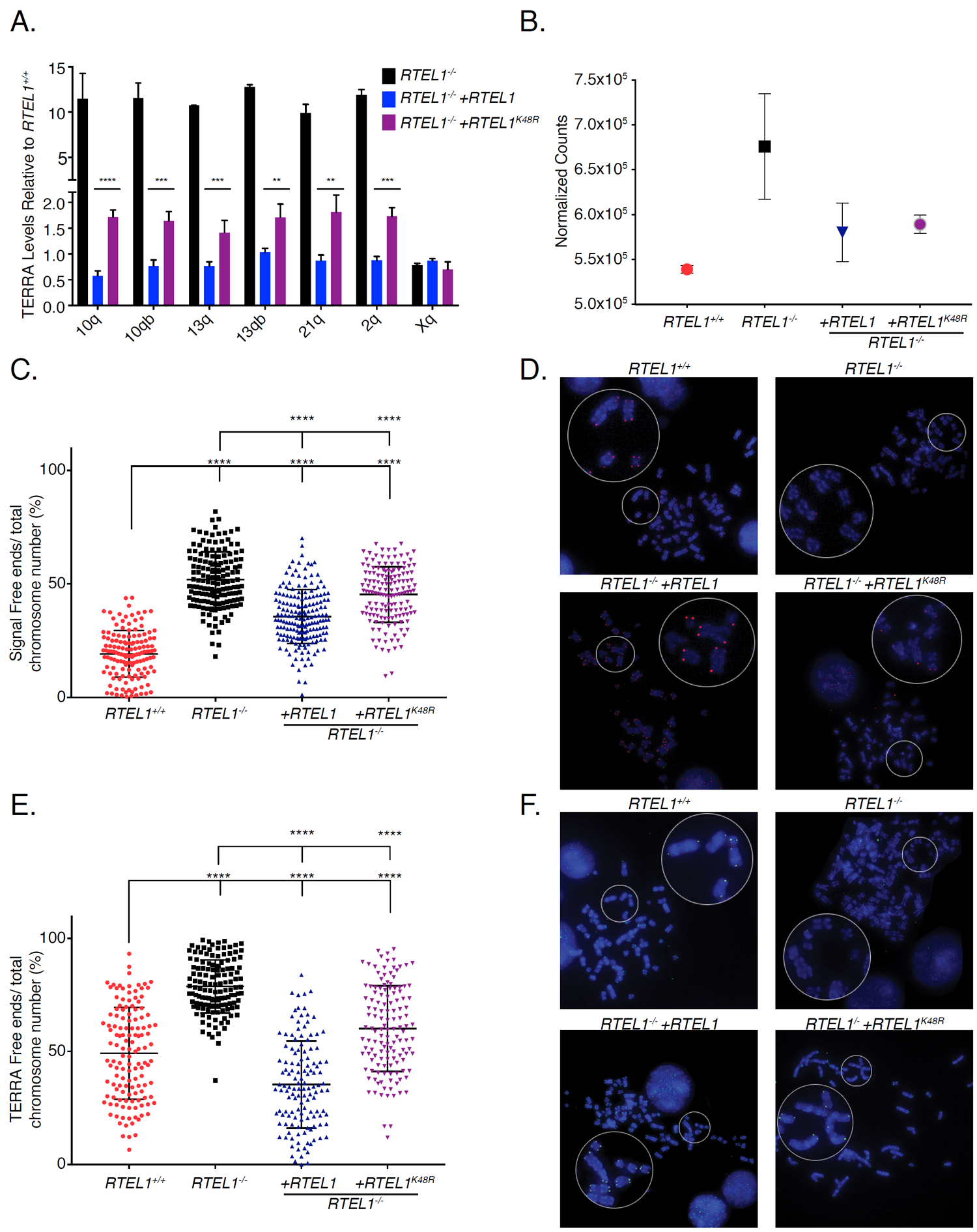
RTEL1 Influences TERRA Localization. TERRA levels are rescued upon RTEL1 complementation. (A) TERRA levels for four upregulated chromosome ends in the RTEL1-KO cell line were tested in cells complemented with *FLAG-RTEL1* and a catalytic site mutant *FLAG-RTEL1*^*K48R*^. Bars represent the relative TERRA expression measured by qRT-PCR in three independent experiments (*p < 0.05, **p <0.01, ***p < 0.001, t-test). (B) Total subtelomeric counts determined by RNA-seq experiments in the cell lines specified in (A). Mean +/- SEM is shown. (C) Quantification of telomere loss per chromosome in *FLAG-RTEL1* and *FLAG-RTEL1*^*K48R*^ reconstituted cell lines by telomere FISH. (D) Representative telomere FISH images are shown with zoomed-in section (large white circle for zoom of small white circle). (E) Quantification of TERRA loss per chromosome in *FLAG-RTEL1* and *FLAG-RTEL1*^*K48R*^ reconstituted cell lines by TERRA FISH. (F) Representative TERRA FISH images are shown with zoomed-in section (large white circle for zoom of small white circle). Data for both TERRA and Telomere FISH experiments represent the average of at least 50 metaphases in each of three independent experiments and three independent reconstitutions shown as the mean and standard deviation (*p < 0.05, **p <0.01, ***p < 0.001, t-test). (See also Figure S7).

The normalization of TERRA levels in complemented cells was not correlated with restoration of telomeric DNA. We examined the levels of telomere loss in RTEL1-KO cells and cells complemented with *FLAG-RTEL1* and *FLAG-RTEL1*^*K48R*^ by Tel FISH. The frequency of telomeric signal free ends was only partially reduced in the reconstituted samples: 19%, 52%, 36%, and 45% for RTEL1 wild type, RTEL1-KO, *FLAG-RTEL1*, and *FLAG-RTEL1*^*K48R*^ reconstitutions respectively (**Figure 4C and 4D**).

### RTEL1 Influences TERRA Localization

We next examined TERRA localization in RTEL1-KO cells. TERRA is localized to telomeric regions within R-loop structures that are established co-transcriptionally or post-transcriptionally by insertion of TERRA in trans (Feretzaki et al., 2019). To assess whether TERRA’s telomeric localization was affected by RTEL1 deficiency, we carried out RNA FISH for TERRA on metaphase spreads of RTEL-KO cells and cells complemented with *FLAG-RTEL1* and *FLAG-RTEL1*^*K48R*^ by TERRA FISH (**Figure 4E and 4F**). The level of TERRA free telomeres in wild type cells was 49.2%, while the majority of the RTEL1-KO chromatids (78.8%) had no detectable TERRA signal. Complementation with wild type *RTEL1* reduced the number of TERRA free ends significantly, with the *FLAG-RTEL1* having a greater impact than *FLAG-RTEL1*^*K48R*^ (35.4% *vs*. 60% TERRA free ends). These data offer a novel perspective on the role of RTEL1 and TERRA in stabilizing telomeric ends, suggesting that RTEL1 has a role in the establishment or stability of TERRA containing R-loops while at the same time influencing the steady state levels of TERRA RNA.

These data raise the possibility of a relationship between TERRA R-loops and TERRA levels. Therefore, we targeted R-loops for destruction by overexpressing RNAse H1. *RTEL1* wild type cells and *RTEL-KO*-*FLAG-RTEL1-* complemented cells were transfected with *RNAse H1-GFP*. Following 12 days of G418 selection, the selected pool of cells was assayed for RNAse H1-GFP expression (**Figure 5A**), and the levels of and localization of TERRA were assessed. We found that TERRA levels in the RNAse H1 expressing lines were significantly elevated, thus partially phenocopying *RTEL1* deficiency (**Figure 5B, Figure 3D, and S8A**).

**Figure 5.**
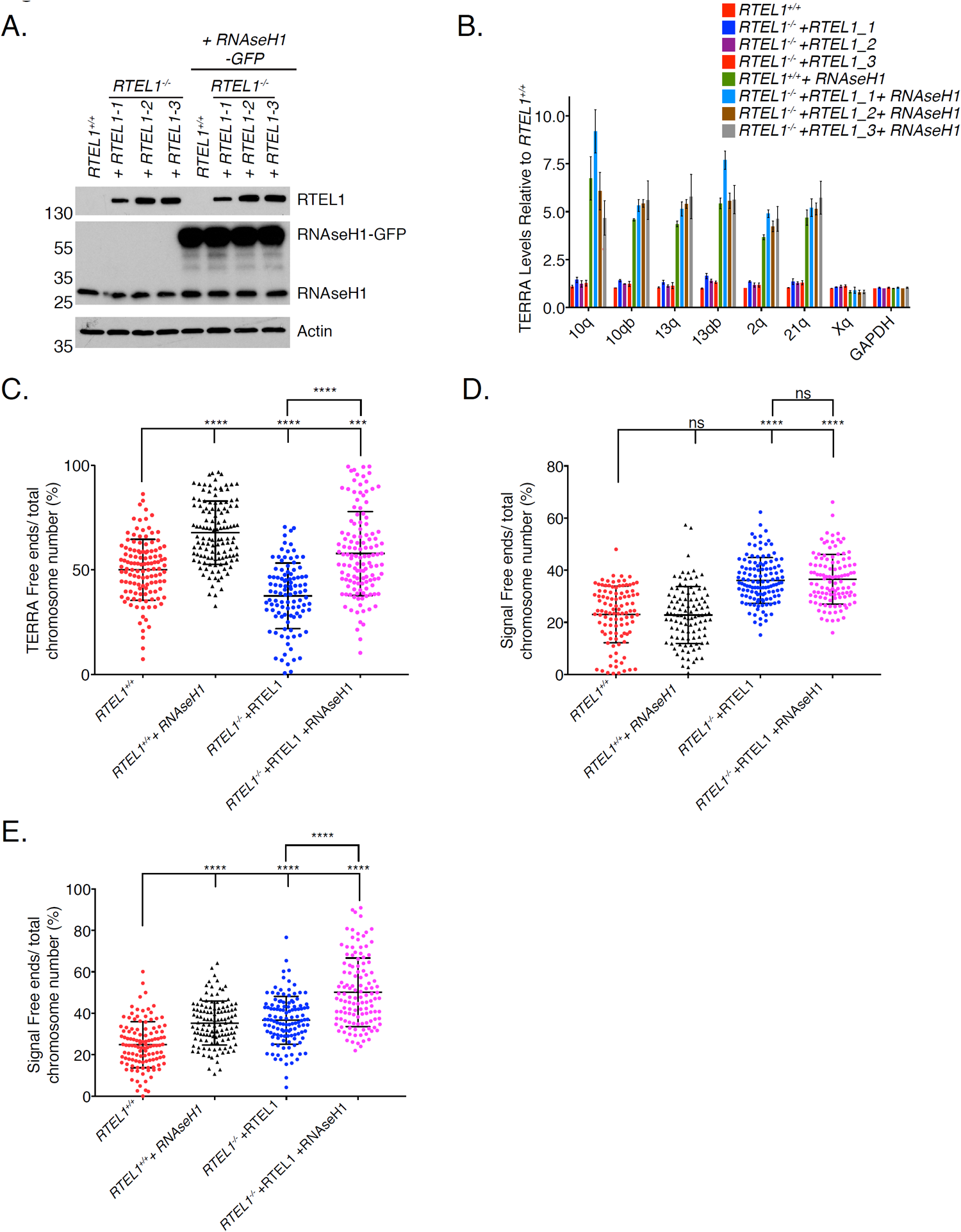
*RNAse H1* overexpression Partially Phenocopies RTEL1 Deficiency. TERRA levels are elevated after RNAse H1 overexpression. (A) Western blotting of wild type HEK293 cells and RTEL1-KO cells complemented with *FLAG-RTEL1* and transfected with *RNAse H1-GFP*. (B) TERRA levels of specified chromosome ends were tested in cells transfected with *RNAse H1-GFP*. Bars represent the relative TERRA expression measured by qRT-PCR in three independent experiments (*p < 0.05, **p <0.01, ***p < 0.001, t-test). (C) Quantification of TERRA loss per chromosome in cells transfected with *RNAse H1-GFP* by TERRA FISH (22 days in selection). (D) Quantification of telomere loss per chromosome in cells transfected with *RNAse H1-GFP* by telomere FISH (22 days in selection). (E) Quantification of telomere loss per chromosome in cells transfected with *RNAse H1-GFP* by telomere FISH (52 days in selection). Three independent cell lines complemented with RTEL1 were averaged for analysis where specified. FISH experiments represent the average of at least 50 metaphases in each of 2 independent experiments and three independent reconstitutions shown as the mean and standard deviation (*p < 0.05, **p <0.01, ***p < 0.001, t-test). (See also Figure S8).

As expected, the telomeric localization of TERRA was also diminished after *RNAse H1-GFP* overexpression, again partially phenocopying *RTEL1* deficiency (**Figure 5C**). The level of TERRA free ends in the wild type cells were 50%, while 68% of wild type RTEL1 cells with *RNAse H1-GFP* had no TERRA signal. Similarly, TERRA free ends were increased from 38% to 58% in *FLAG-RTEL1* complemented RTEL-KO cells in which *RNAse H1-GFP* was overexpressed (**Figure 5C**).

The frequency of telomeric signal free ends at 22 days post-transfection was not significantly affected by *RNAse H1-GFP* overexpression: 23%, 23%, 36%, and 37% for RTEL1 wild type, *RTEL1* wild type + *RNAse H1-GFP, RTEL-KO-FLAG-RTEL1* and *RTEL-KO-FLAG-RTEL1+RNAse H1-GFP*, respectively (**Figure 5D**). However, at 52 days, we observed a marked increase in the frequency of signal free telomeric ends; 24.8% *vs*. 35.3% for RTEL1 wild type compared to RTEL1 wild type + *RNAse H1-GFP* and 36.7%, vs. 50.2% in *RTEL-KO-FLAG-RTEL1* and *RTEL-KO-FLAG-RTEL1+RNAse H1-GFP*. In contrast, no significant difference in TERRA levels was observed at this time point (**Figure 5E and S8B**).

## Discussion

In this study, we established that RTEL1 influences the abundance and localization of the telomeric RNA TERRA, likely through direct binding to TERRA via a previously undescribed domain at the RTEL1 C-terminus. *In vitro* analyses revealed that RTEL1 binds G-quadruplex DNA and RNA, but the differential affinity for folded *vs*. unfolded RNA is significantly higher than for the analogous structures in DNA. In cells, *RTEL1* hypomorphism and deficiency are associated with telomere fragility and loss, as noted previously (Ballew et al., 2013a; Le Guen et al., 2013; Sarek et al., 2015; Uringa et al., 2011; Vannier et al., 2012). However, we observed a marked increase in TERRA levels in *RTEL1* deficient cells. Increased levels of TERRA was not associated with increased telomeric localization. Rather, in *RTEL1* deficient cells, the frequency of TERRA occupancy at the telomeres of metaphase chromosomes was markedly reduced.

Reconstitution of *RTEL-KO* cells revealed that the effects on TERRA were only partially helicase dependent. We also found that *RTEL1* cDNAs encoding RTEL1 variants that lacked the G4 binding domain were unable to support clonal survival of the *RTEL1* knockout line, indicating the physiological importance of G4 engagement by RTEL1. The observation of elevated TERRA levels in *RTEL1* knockout cells was recapitulated in several lymphoblastoid cell lines from HH and DC patients as well as in a fibroblast line derived from the HH patient homozygous for the *RTEL1*^*R1264H*^ allele (Ballew et al., 2013a; Ballew et al., 2013b). In each of those cases, *RTEL1*^*1010X*^, *RTEL1*^*R1264H*^, and the *RTEL1*^*c*.*2142-7C>G*^ mutations predicted to alter splicing (Ungar et al., 2018), the RTEL1 C-terminus is affected (**Figure 1A, 3E and 3F**). These data raise the possibility that telomere dysfunction may be caused by alterations of TERRA abundance and localization associated with RTEL1 deficiency, and this may contribute to the clinical phenotypes of DC and HH.

In addition to binding TERRA, the RTEL1 C-terminal domain promotes oligomerization via the C4 domain (**Figure 1C**). In this regard, it is notable that some *RTEL1* alleles behave as autosomal dominant mutations with respect to clinical presentation (*RTEL1*^*1010X*^) or sub-clinically with respect to telomere length; the *RTEL1*^*R1264H*^ heterozygous parents of homozygous *RTEL1*^*R1264H*^ patients exhibit pronounced telomere shortening (Ballew et al., 2013a; Ballew et al., 2013b; Ungar et al., 2018). Supporting the importance of oligomerization, overexpression of *RTEL1*^*R1264H*^ in wild type HEK293 cells leads to rapid selection against *RTEL1*^*R1264H*^ expressing cells, whereas overexpression of *RTEL1* cDNAs that encode proteins lacking the RING domain was well tolerated (data not shown). Since the RTEL1 helicase domain is an independent dimerization interface, the apparent toxicity of the RTEL1^R1264H^ may reflect that RTEL1 assemblies in which the C-terminal homotypic interaction is impaired are toxic. The RING domain specifies functions aside from dimerization. For example, the interaction of RTEL1 with TRF2 is lost in the *RTEL1*^*R1264H*^ mutant (Sarek et al., 2016); hence the apparent toxicity of *RTEL1*^*R1264H*^ may not be due to an effect on dimerization. Nevertheless, these data support a model where both oligomerization and RNA binding are important for RTEL1 function independent of its helicase activity.

On the basis of the data obtained, we propose that the formation and maintenance of the TERRA R-loop at the telomere influences TERRA transcription and promotes telomere stability. In this conception, RTEL1 would be involved in either facilitating the creation of the TERRA containing telomeric R-loop or contributing to its maintenance. Once established, the TERRA R-loop would limit further transcription from R-loop containing chromosome ends (**Figure 6**). This part of the model is based on the reduction in TERRA R-loops and increased levels of TERRA in RTEL1 deficient cells (**Figure 3C-3F and 4E-4F**). It is likely that the ability of RTEL1 to bind and discriminate distinct nucleic acid folds *in vitro* is biologically relevant at the telomere where these structures are predicted to form. Implicitly, the model also predicts that the presence of the R-loop at the telomere contributes to telomere stability.

**Figure 6.**
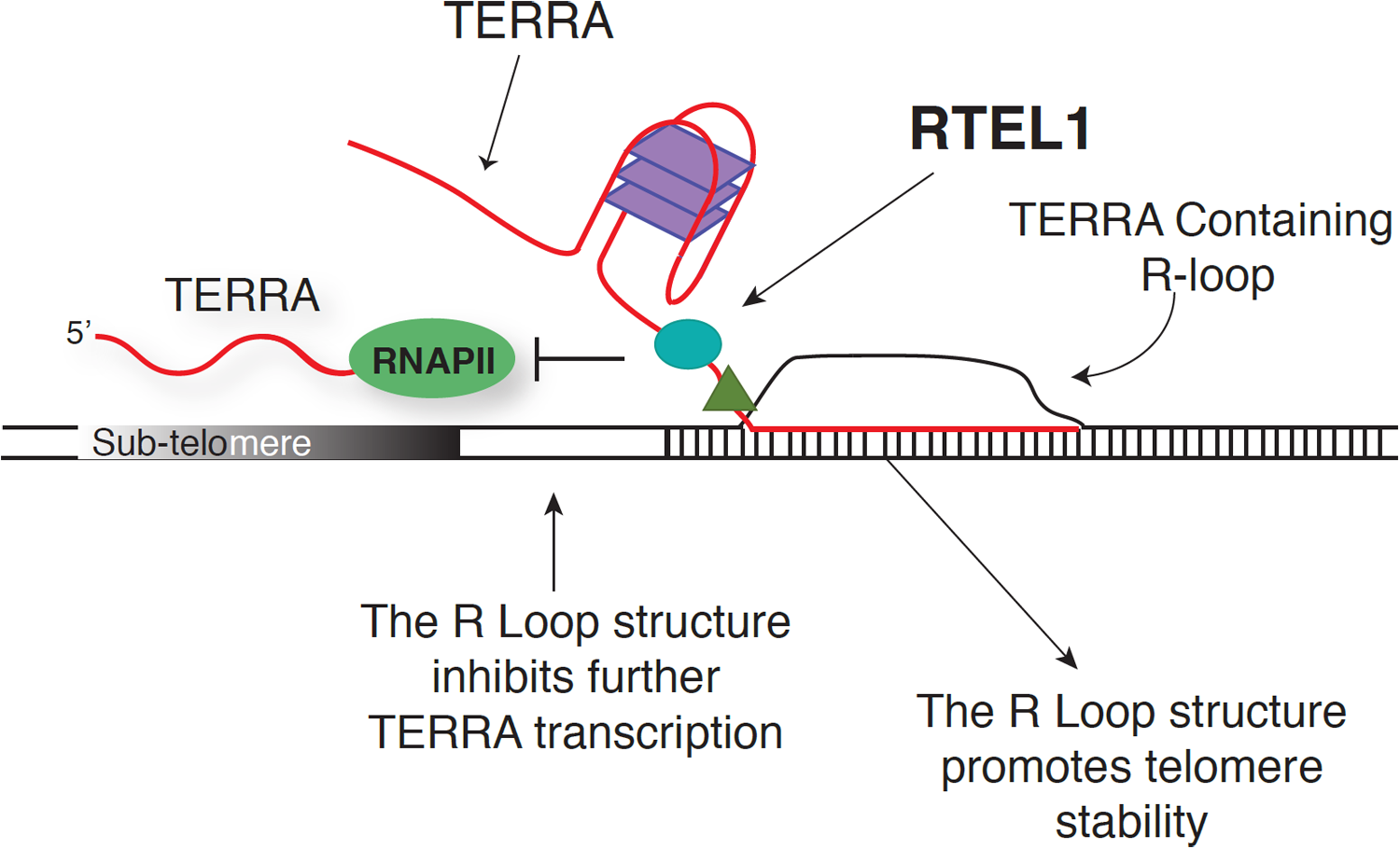
RTEL1 Influences the Abundance and Localization of the Telomeric RNA TERRA. The formation and maintenance of the TERRA R-loop at the telomere influences TERRA transcription and promotes telomere stability. We propose RTEL1 would be involved in either facilitating the creation or contributing to the maintenance of the TERRA containing telomeric R-loop. Once established, the TERRA R-loop would limit further transcription from R-loop containing chromosome ends

The phenotypic outcomes of *RNAse H1* overexpression, which is known to disrupt R-loops (Arora et al., 2014), were consistent with the model proposed. First, *RNAse H1* overexpression increased the frequency of metaphase chromosomes that lacked TERRA FISH signal. This apparent reduction in TERRA R-loops was associated with increased levels of TERRA RNA, as inferred by Q-PCR. Notably, *RNAse H1* overexpression increased TERRA levels from chromosomes 2q, 10q, 13q, and 21q (**Figure 5B**), but not chromosomes 8p, 19p or Xq (**Figure 5B**, and data not shown). Transcription of TERRA from those same chromosomes also increases upon *RTEL1* inactivation and in hypomorphic patient cells (Figure 3D-F, S7C and S7D), strengthening the idea that the R-loop exerts an influence on TERRA abundance, at least from those telomeric loci.

Second, the model predicts that the TERRA R-loop would contribute to telomere stability, and accordingly that *RNAse H1* overexpression would decrease telomere stability. At 10 and 22 days after transfection of *RNAse H1*, R-loops, as inferred from TERRA RNA FISH, were diminished but no increase in telomere-free chromatid ends was evident, consistent with previous data ((Arora et al., 2014) and **Figure 5C and 5D**). As noted above, TERRA levels were elevated at those time points. At 52 days post transfection, a significant increase in telomere loss was evident (**Figure 5E**), again consistent with the model.

However, at the 52 day time point, TERRA levels were no longer elevated. On its face, this observation is inconsistent with the simplest interpretation of the model, but it is conceivable that prolonged *RNAse H1* overexpression may exert an indirect effect on TERRA expression. R-loops have been shown to prevent methylation of promoters by excluding binding of a methyltransferase required for proper gene silencing (Ginno et al., 2012). Some TERRA promoters are regulated by DNA methylation and depletion of, or defects in methyltransferases lead to elevated TERRA levels (Feretzaki et al., 2019; Nergadze et al., 2009; Sagie et al., 2017). TERRA also interacts with ATRX, and antagonizes its functions (Chu, Cifuentes-Rojas et al. 2017). Hence it is also conceivable that prolonged absence of TERRA-containing R-loops allows ATRX-mediated silencing of TERRA promoters. TERRA expression may also be reduced in an R-loop independent manner.

The role of RTEL1 in regulating R-loops is likely to be complex, and may involve both negative and positive regulation of those structures, likely through its binding to G4 structures. R-loops are associated with GC rich genomic contexts (Sanz et al., 2016) and recent data suggest that RTEL1 dismantles R-loops at loci in which G-quadruplex forming sequences are common, including those within common fragile sites (Wu, 2020).

On the other hand, there is precedent for G4 binding helicases promoting the post-transcriptional formation of R-loop structures. For example, immunoglobulin class switch regions are G-rich repetitive elements that must be transcribed in order for class switch recombination to occur. Debranching of the spliced switch region transcript and its subsequent insertion into an R-loop structure requires the action of DDX1, an RNA helicase that binds the G4 structure of the transcript and inserts the RNA into the switch region DNA duplex to form the R-loop structure required for class switch recombination (Ribeiro de Almeida et al., 2018; Yewdell and Chaudhuri, 2017). In addition, it is clear that TERRA can be inserted into R-loop structures in trans (Chu et al., 2017; Lee et al., 2018), although the helicase required and whether it depends on G4 structure is not known.

The model proposed here wherein RTEL1 promotes telomere stability via its influence on the localization and abundance of TERRA RNA does not exclude other models to explain the role of RTEL1 in stabilizing the telomere. A prevalent model suggests that RTEL1 disassembles T-loops to allow DNA replication to proceed to the chromosome end (Sarek et al., 2015; Vannier et al., 2012). When RTEL1 is depleted the T-loops are persistent and inappropriately resolved by the SLX4 nuclease. Loss of RTEL1-dependent resolution of the T-loop or other secondary structures has also been suggested to promote replication fork reversal upstream of the secondary structure obstacle. This pathological situation is proposed to be stabilized by binding of the TERT protein to the reversed fork and again lead to cleavage by SLX4 (Margalef et al., 2018).

These models account for the detection of extrachromosomal telomeric circles, and for telomere fragility. A prediction of these models is that telomere loss should be identical on both sister chromatids, since the SLX4 cleavage event occurs before replication. Based on telomere FISH, this is true for a subset of chromosomes, but loss of telomeric signal on one sister chromatid is often observed ((Ballew et al., 2013a; Sarek et al., 2015; Vannier et al., 2012) and **Figure 4D** herein). This signal heterogeneity suggests that there may be more than one mechanism for how the telomeres are maintained by RTEL1, with some loss of telomeric sequences arising via the model proposed here (**Figure 6**). The data presented herein support a novel mechanism to account for RTEL1’s influence on telomere stability, and may illuminate the biological significance of RTEL1-TERRA interaction. The data support the view that TERRA plays a role in stabilizing telomeric sequences.

## Acknowledgements

We thank Christopher Lima and John Zinder for valuable advice and technical assistance throughout this project. We thank Dinshaw Patel and Wei Xie for help with SEC-MALS, and the members of the MSKCC proteomics core for assistance with proteomics analysis. We are grateful to all the members of the Petrini laboratory and Tom Kelly for helpful discussions. We thank Laura Feeney and Tom Kelly for critical reading of the manuscript. This work was supported by GM56888 (J.H.J.P), U54 OD020355 (J.H.J.P), the MSK Cancer Center Core Grant P30 CA008748 (J.H.J.P), the DOD award W81XWH-16-1-0218 (F.G), and the intramural research program of the Division of Cancer Epidemiology and Genetics, National Cancer Institute (S.A.S).

## Author Contributions

F.G, A.G, H.W, C.C, and M.H performed experiments. F.G, A.G, H.W, M.H, and J.H.J.P analyzed data. S.A.S evaluated patients and contributed the patient lymphoblastoid cell lines. F.G and J.H.J.P wrote the manuscript. J.H.J.P and T.T provided general supervision and mentorship.

## Materials and Methods

### Cell lines

HEK293 cells were grown in 10% DMEM and 10% FBS. EBV-immortalized lymphoblastoid cell lines were derived from participants in the National Cancer Institute’s IRB-approved longitudinal cohort study entitled “Etiologic Investigation of Cancer Susceptibility in Inherited Bone Marrow Failure Syndromes” (ClinicalTrials.gov Identifier: NCT00027274) (Alter et al., 2018). This study includes comprehensive family history and individual history questionnaires, detailed medical record review, and biospecimen collection. Detailed clinical evaluations were performed at the NIH Clinical Center per protocol. LCLs were grown in RPMI Media supplemented with 100 U/ml penicillin, 100 μg/ml streptomycin, GlutaMAX (Life Technologies), and 20% FBS. hTERT transduced fibroblasts were cultured in DMEM media supplemented with 100 U/ml penicillin, 100 μg/ml streptomycin, GlutaMAX, and 15% FBS. All cell lines used were routinely checked for mycoplasma.

### Protein Expression and Purification

Sequences encoding residues 762-1300, 762-1097, 1097-1300, 1118-1300, and 1143-1300 of human RTEL1 were amplified by PCR then subcloned between the *EcoRI* and *XhoI* restriction sites of *pET28* (Novagen). RTEL1 proteins were expressed as 6-histidine-fusions in the *E. coli* strain BL21 and purified using His-trap (GE Healthcare), Hi-Trap Q (GE Healthcare), and Superdex 200 columns (GE Healthcare). For expression in mammalian cells, full length human *RTEL1* and *RTEL1*^*1-1097*^ were subcloned in to *pcDNA3*.*1 MCS-BirA (R118G)-HA* between *HpaI* and *EcoR1* sites. *pcDNA3*.*1 MCS-BirA(R118G)-HA* was a gift from Kyle Roux (Addgene plasmid # 36047;http://n2t.net/addgene:36047; RRID:Addgene_36047). Full length *RTEL1* was cloned into *pDest12*.*2-myc-his* and *pDESTb-FRT-C3xFLAG*. Residues 1097-1300 were amplified by PCR and subcloned between *EcoRI* and *XhoI* restriction sites of *pCDNA3*.*1_FLAG-HA* (Invitrogen) and *pIC113* (*pIC113* was a gift from Iain Cheeseman and Arshad Desai (Addgene plasmid # 44434; http://n2t.net/addgene:44434; RRID: Addgene_44434)). An additional nuclear localization signal (MADPKKKRK) was added to the 5’ end of the *RTEL1* truncation constructs and mutagenesis was performed using standard protocols. For expression of RTEL1 proteins in HEK293 RTEL1-KO cells, full length human *RTEL1* was cloned into *pLenti-C-myc-DDK-IRES-Puro* (Origene) plasmid by digestion with *AscI* and *MIuI* and mutagenesis for *RTEL1*^*K48R*^ was performed using QuikChange Lightning Multi Site-Directed Mutagenesis Kit (Agilent). Cells were infected with lentiviruses expressing the indicated RTEL1 proteins and infected cells were selected with puromycin as previously described (Garzia et al., 2017a). Cell lines derived from three independent infections were used for reconstitution experiments. The *pEGFP-RNAse H1* was a gift from Andrew Jackson & Martin Reijns (Addgene plasmid # 108699; http://n2t.net/addgene:108699; RRID: Addgene_108699). HEK293 were transfected using PEI-25K (polysciences Inc.) and after 48 hours selected with G418 for the indicated time points.

### Co-immunoprecipitation and Western Blotting

HEK293T cells were transfected with wild type and single point mutants of the following constructs as indicated: *RTEL1-myc-HIS, RTEL1-FLAG, GFP-RTEL1*^*Δ1097*,^ and *FLAG-HA RTEL1*^*Δ1097*^ using PEI-25K (polysciences Inc.). Cell extracts were prepared using 1X PBS, 0.5% (v/v) TritonX-100 and protease inhibitor tablet (Sigma). Proteins were immunoprecipitated by incubating cell extracts overnight with either FLAG M2 (Sigma) or GFP (Clonetech) antibodies at 4°C, followed by incubation with Protein A/G beads for 3 hours. Lentiviral transfections were performed as previously described (Garzia et al., 2017b). Western blotting was performed using standard methods. Total extracts were prepared in lysis buffer (60 mM Tris-HCl pH 6.8, 2% SDS) and analyzed with specified antibodies. The antibodies used in this study were GFP JL-8 (Clonetech, 632381), ANTI-FLAG® M2 (Sigma, F1804), HA (Biolegend #901515), myc 9E10 (BioXcell, BE0238), RTEL1 (custom made), Actin-HRP (ABCAM, ab49900), GAPDH (Santacruz, sc_32233) and RNAse H1 (Proteintech 15606-1-AP).

### SEC-MALS

The molar masses of RTEL1 proteins were analyzed using Size exclusion chromatography coupled to multi-angle light scattering (SEC-MALS). 200-300 *µ*g of the indicated proteins were injected into a Superdex 200 10/300 GL (GE Healthcare) equilibrated at a flow rate of 0.3 ml per minute in 50 mM Hepes pH 7.5, 1 mM TCEP, and either 150 or 300 mM NaCl. Light scattering was monitored with a miniDAWNTREOS system (Wyatt Technology), concentration was measured with the Optilab T-rEX differential refractometer (Wyatt Technology), and molar masses were calculated using the Astra 6.1 program (Wyatt Technology) with a dn/dc value of 0.185 ml/g.

### BioID

Affinity purification of biotinylated proteins was performed as previously described (Roux et al., 2012). Briefly cells were supplemented with 50 *µ*M biotin for 24 hours and sonicated in lysis buffer containing 50 mM Tris, pH 7.4, 500 mM NaCl, 0.4% SDS, 5 mM EDTA, 1 mM DTT, 1x Complete protease inhibitor (Roche). Triton X-100 was brought up to 2% and an equal volume of 50 mM Tris, pH 7.4 was added before clarifying the lysate by centrifugation. Supernatants were incubated with 200 *µ*l Dynabeads (MyOne Steptavadin C1; Invitrogen) overnight. The beads were washed with 1.5 ml wash buffer 1 (2% SDS), 1.5 wash buffer 2 (0.1% deoxycholate, 1% Triton X-100, 500 mM NaCl, 1 mM EDTA, and 50 mM Hepes, pH 7.5), 1.5 ml wash buffer 3 (250 mM LiCl, 0.5% NP-40, 0.5% deoxycholate, 1 mM EDTA, and 10 mM Tris, pH 8.1). After the wash 1.5 ml (50 mM Tris, pH 7.4, and 50 mM NaCl) and 50 *µ*l 50 mM NH_4_HCO_3_ were added before mass spectrometry analysis. On-bead tryptic digestions were analyzed by the MSKCC Proteomics Core as previously described (Roux et al., 2012). MassHunter Qualitative Analysis software (Agilent Technologies) was used to analyze the raw data and database search was done using MASCOT (MatrixScience). SCAFFOLD Q+ (Proteome Software) was used to visualize the data with a threshold of at least 2 identified peptides and a minimum 95% probability each.

### Binding Assays

Oligonucleotides were refolded in 10 mM Tris pH 7.5 and 100 mM KCL (unless otherwise noted) with slow cooling starting at 95°C down to room temperature. The change in fluorescence anisotropy of the indicated 5’ fluorescein labeled oligonucleotides was measured to determine the relative binding affinities of RTEL1 proteins. The labeled oligonucleotides were mixed in 20 μl reaction volumes with the indicated RTEL1 proteins at concentrations ranging from 0-5 μM (unless otherwise indicated) in a reaction buffer containing 20 mM Tris, pH 7.5, 50 mM KCl, 0.5 mM TCEP, 0.5 mM MgCl2, 10% glycerol, and 0.1% IGEPAL in a 384-well microplate. The binding data were collected on a SpectraMax M5 (Molecular Devices) using a 495 nm excitation wavelength and a 525 nm emission wavelength. Apparent K_d_ values were calculated from triplicate experiments using a model for receptor depletion and plotted using Prism 7 GraphPad Software. The model used was:

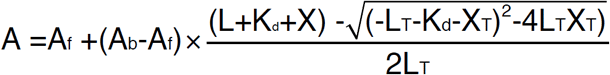

Where A is the anisotropy measured; A_b_ is the anisotropy at saturation (100%), A_f_ is the anisotropy of the free oligonucleotide, L is the fixed concentration of the oligonucleotide and X is the protein concentration. For competition assays, RTEL1-FAM-TERRA-MUT (wild type TERRA RNA was not readily competed by any of the oligonucleotides tested) complex concentration was maintained at 200 nM or 1 *µ*M as indicated. Competitor oligonucleotides were added at concentrations ranging from 0-20 *µ*M as specified. Experiments were performed in triplicate, and the data were fit to a one-site binding model for competition assays using Prism 7. For EMSA (Electrophoretic mobility shift assay), binding reactions were performed in buffer containing 20 mM Tris, pH 7.5, 50 mM KCl, 0.5 mM TCEP, 0.5 mM MgCl2, 10% glycerol, and 0.1% IGEPAL. RTEL1 proteins and 5’ FAM-labeled oligonucleotide at the indicated concentrations were incubated at room temperature for 30 minutes then loaded onto a 4-20% polyacrylamide non-denaturing gel. The gels were imaged using fluorescein fluorescence on a Typhoon FLA9500 instrument (GE).

### RT-qPCR

Total RNA was extracted from the specified cell lines using the RNeasy Mini Kit (Qiagen) with DNase treatment. cDNA was synthesized using Superscript IV kit (Invitrogen) using random primers. TERRA expression in each sample was normalized to *GAPDH* and relative expression was determined with the comparative CT method. Primers used in this study are listed in Supplementary Table 1.

**Table 1.**
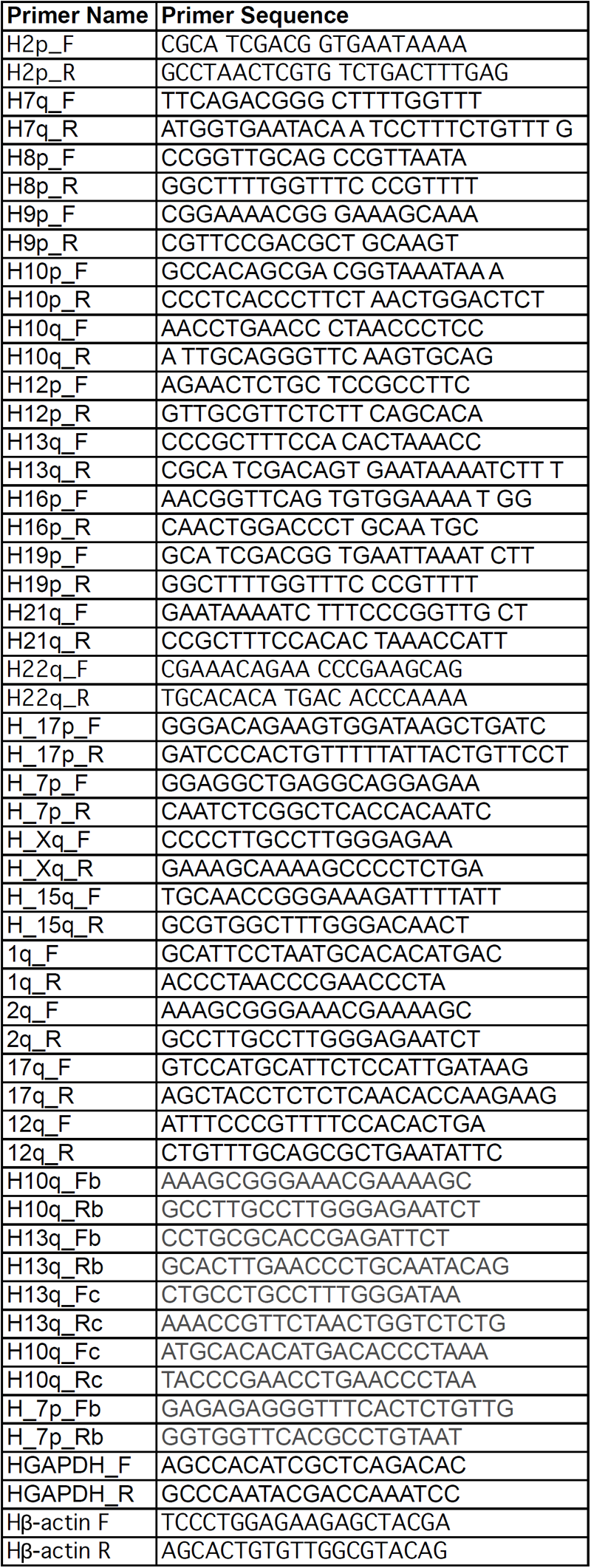
qRT-PCR Primers used in this study.

### DNA and RNA FISH

Metaphase chromosome spreads were prepared by incubating cells with 0.1 *µ*g/*µ*l KaryoMAX colcemid solution for 90 minutes. Cells were harvested at 1000 rpm and resuspended in 0.075 M KCL at 37°C for 15 minutes. Cells were fixed in a 3:1 mixture of ice-cold methanol/acetic acid at least overnight and spread onto glass slides then air dried overnight. Samples were treated with 100 *µ*g/ml RNAse A for 1hr at 37°C, dehydrated through a series of ethanol washes at 70%, 90% and 100% ethanol for 5 minutes each at room temperature, then allowed to air dry. Hybridization was in buffer containing 10 mM Tris pH 7.5, 70% formamide, 0.5% blocking reagent (Roche) and 0.5 *µ*g/ml CY-3 (CCCTAA)_3_ (PNA BIO), denatured at 75°C for 5 minutes and incubated at room temperature for 16hr. Slides were washed twice in buffer containing 10 mM Tris pH 7.5, 0.1% BSA, and 70% formamide, then three times in PBS/0.15% Triton X-100. The slides were mounted using proLong Gold antifade reagent with DAPI (Invitrogen). For TERRA fish slides were treated with a mixture of RNAse A and RNAse H1 for 16 hours or PBS and treated as above with the exception of the denaturation step. Images were acquired on a Deltavision Imaging Elite System (GE Healthcare) with a CMOS Camera on an Olympus IX-71 microscope using a 60X objective. Images were analyzed using ImageJ and statistical analysis was done with Prisim 7 (Graphpad).

### Generation of the *RTEL1* Knockout Line

RTEL1 knockout was performed using CRISPR-Cas9-mediated genome editing as described previously (Ran et al., 2013). We used a guide RNA mapping to Exon 2 of the *RTEL1* gene 5′-TGCCCGCAAGATTGCCGAGA. PCR-genotyping was performed using the following primers: 5′-GGGACTTGCCTGTGGACTTCTCCGC and 5′-CGCCATCCCTTGCCAACCATCCCC. PCR products were cloned using the Zero Blunt PCR Cloning Kit (Thermo Fisher Scientific, K270040) and 20 clones were sequenced per cell line to verify successful genome editing.

### RNA Crosslinking Experiment

To analyze the RNA crosslinking capacity of wild type and mutant RTEL1 proteins, RTEL1-KO cells were complemented with N-terminally FLAG-HA-tagged wild type or mutant RTEL1. Cells were fed with 4-thiouridine (100 M) for 16 hours and crosslinked at 365 nm. A modified version of PAR-CLIP was performed (Garzia, Meyer et al. 2017), using a single RNase A digestion step and using anti-FLAG-M2 magnetic beads (Sigma, M8823) for IP.

### RNA-Seq Analysis for Subtelomere Counts

RNAseq libraries were prepared from total RNA using ribosomal RNA depletion method. Paired-end reads were aligned using STAR against subtelomeric sequences (subtelomeric sequences cover 15 kb before the start of the telomeric sequences TTAGGG for 1p,1q,2p,2q,3p,3q,4p,4q,5p,5q,6p,6q,7p,7q,8p,8q,9p,9q,10p,10q,11q,12p, 12q,13q,14q,15q,16p,16q,17p,17q,18p,18q,19p,19q,10p,10q,21q,22q,Xp,Xq.Yq ends and were provided by Dr. Joachim Linger). Mapped reads were counted using Featurecounts, allowing multi-mapping reads, and counting each alignment fractionally. The raw total counts of all the subtelomeric sequences for each sample were normalized using size factors from DESeq2. The size factors were generated from the whole transcriptome analysis for the same samples.

## Supplementary Figure Legends

**Figure S1.**
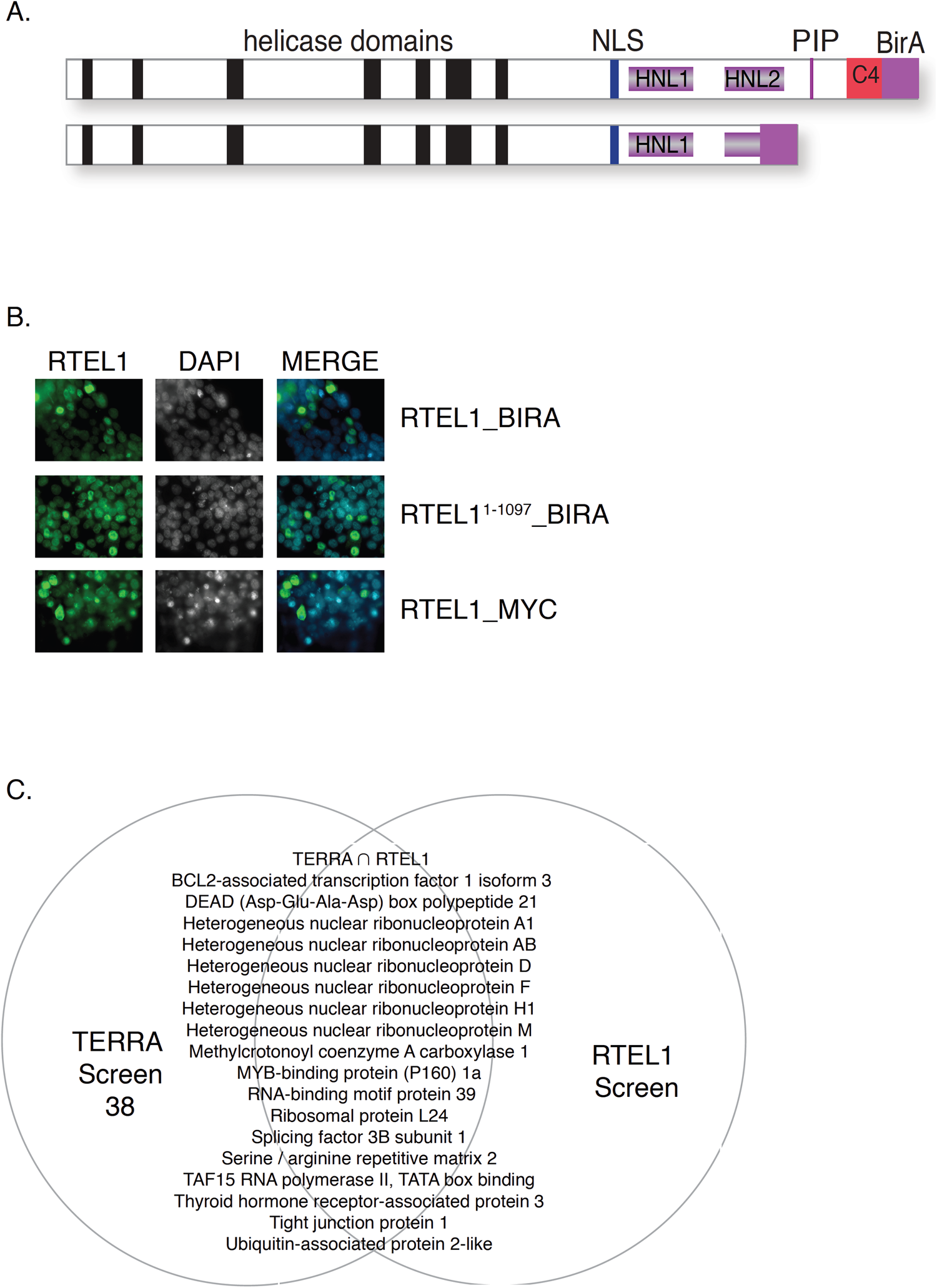
(A) Schematic representation of *RTEL1-BirA* constructs used in this study. (B) RTEL1-BirA fusions did not affect the primarily nuclear localization of RTEL1-BirA compared to myc-tagged RTEL1. (C) Close to 50% of proteins obtained in a Biotin-TERRA pull-down (Lopez de Silanes, Stagno d’Alcontres et al. 2010) matched proteins that were enriched for in the *RTEL1-BirA* construct.

**Figure S2.**
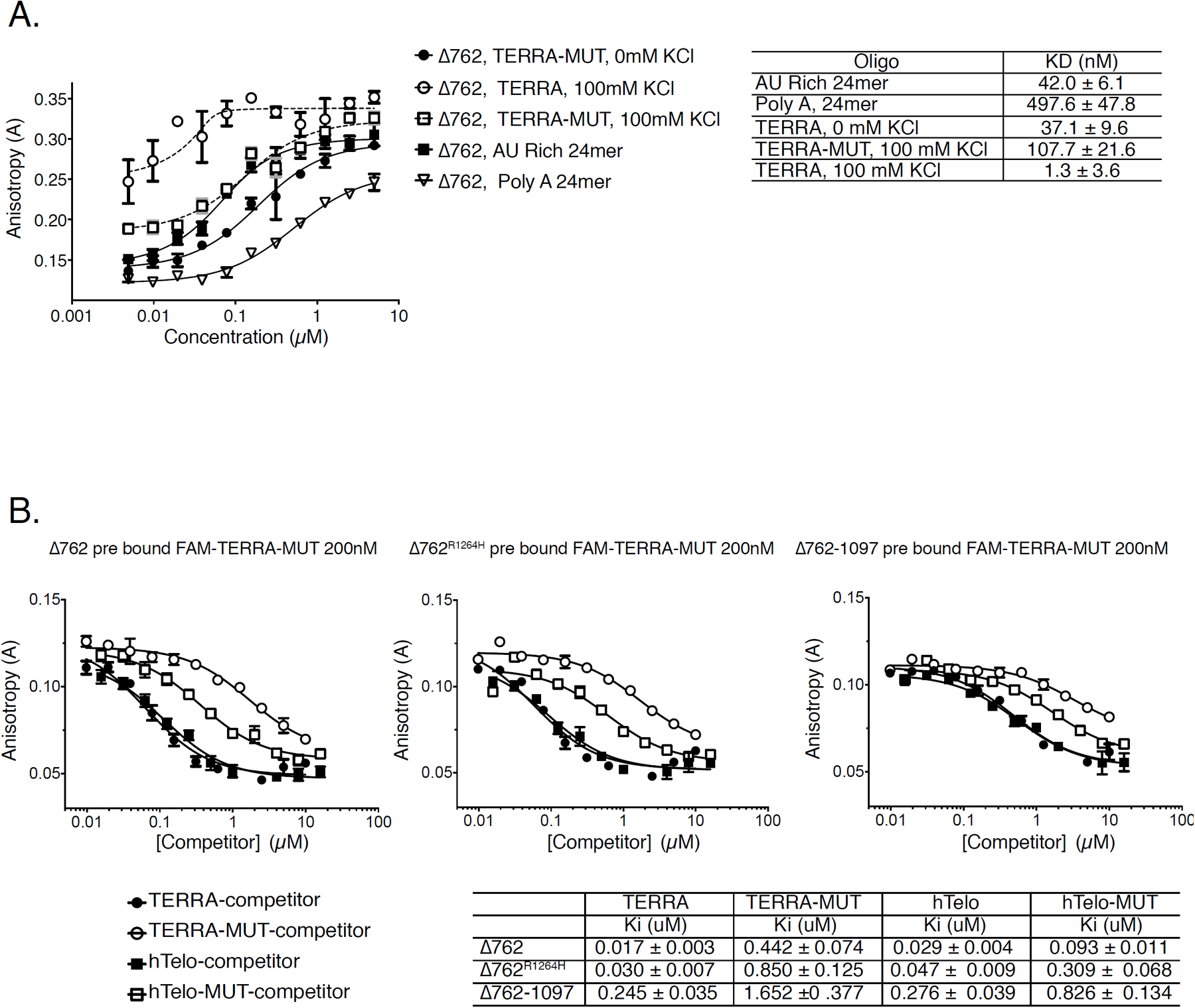
Experimental fluorescence anisotropy data used to calculate apparent dissociation constants for RTEL1 proteins binding to RNA and DNA oligonucleotides indicated in Figure 2. (A) Binding curves for TERRA and TERRA-MUT RNAs folded in the presence and absence of KCl, and AU-rich and polyA RNA controls. (B) Increasing concentrations of the indicated oligonucleotides were added to reactions containing RTEL1 proteins and a 24mer FAM-TERRA-MUT RNA at 200 nM. Curves for the bound FAM-TERRA-MUT competed by the indicated oligonucleotides are shown. All binding assays were conducted in triplicate and mean and standard deviation are shown.

**Figure S3.**
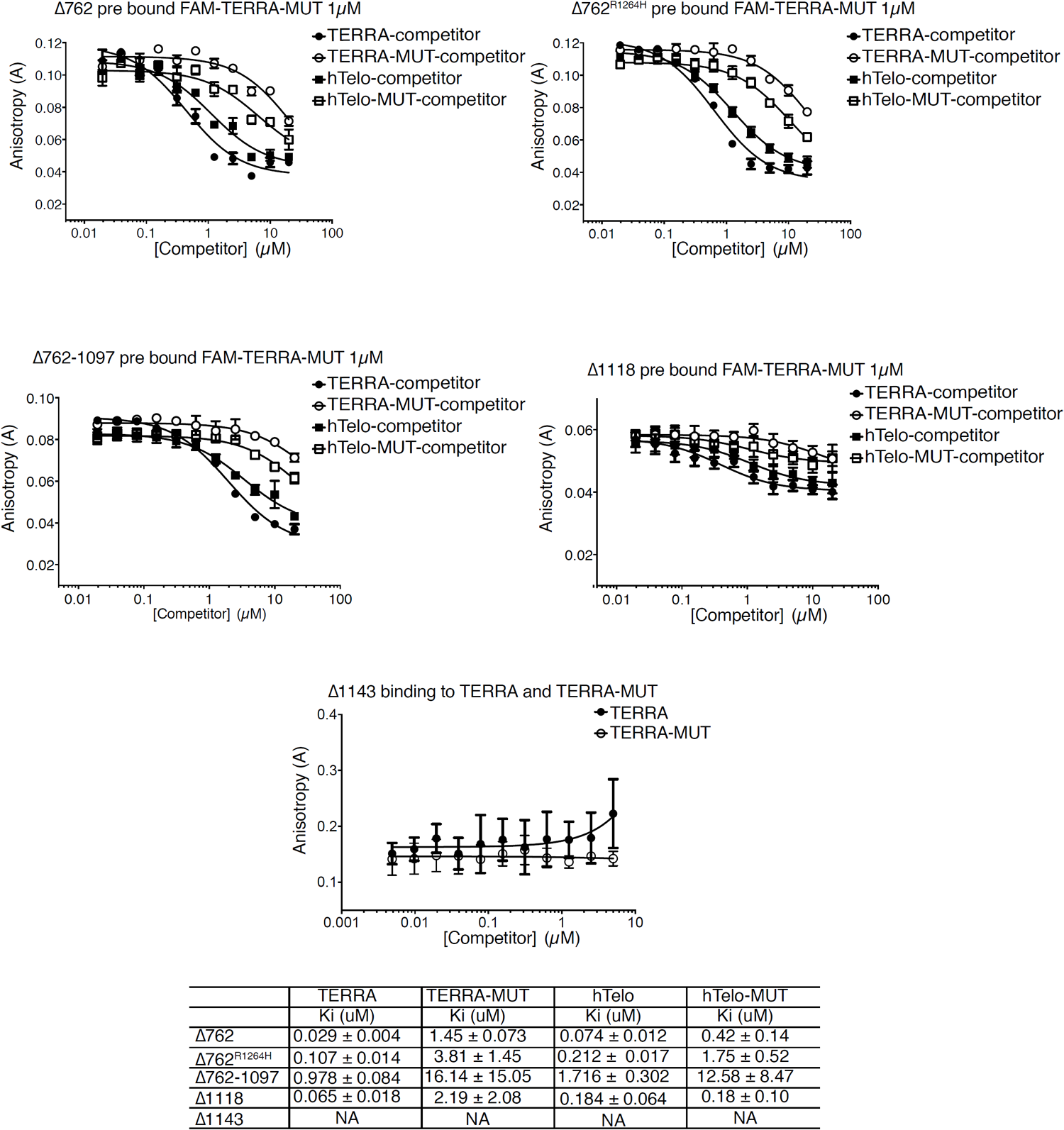
Experimental fluorescence anisotropy data used to calculate apparent dissociation constants for RTEL1 deletion constructs binding to RNA and DNA oligonucleotides indicated in Figure 2. Increasing concentrations of the indicated oligonucleotides were added to reactions containing the indicated RTEL1 proteins and a 24mer FAM-TERRA-MUT RNA at 1 *µ*M. Binding curves used to calculate apparent dissociation constants (Ki) are shown. For RTEL^Δ1143^ direct binding to TERRA and TERRA-MUT RNAs is shown. All binding assays were conducted in triplicate and mean and standard deviation are shown.

**Figure S4.**
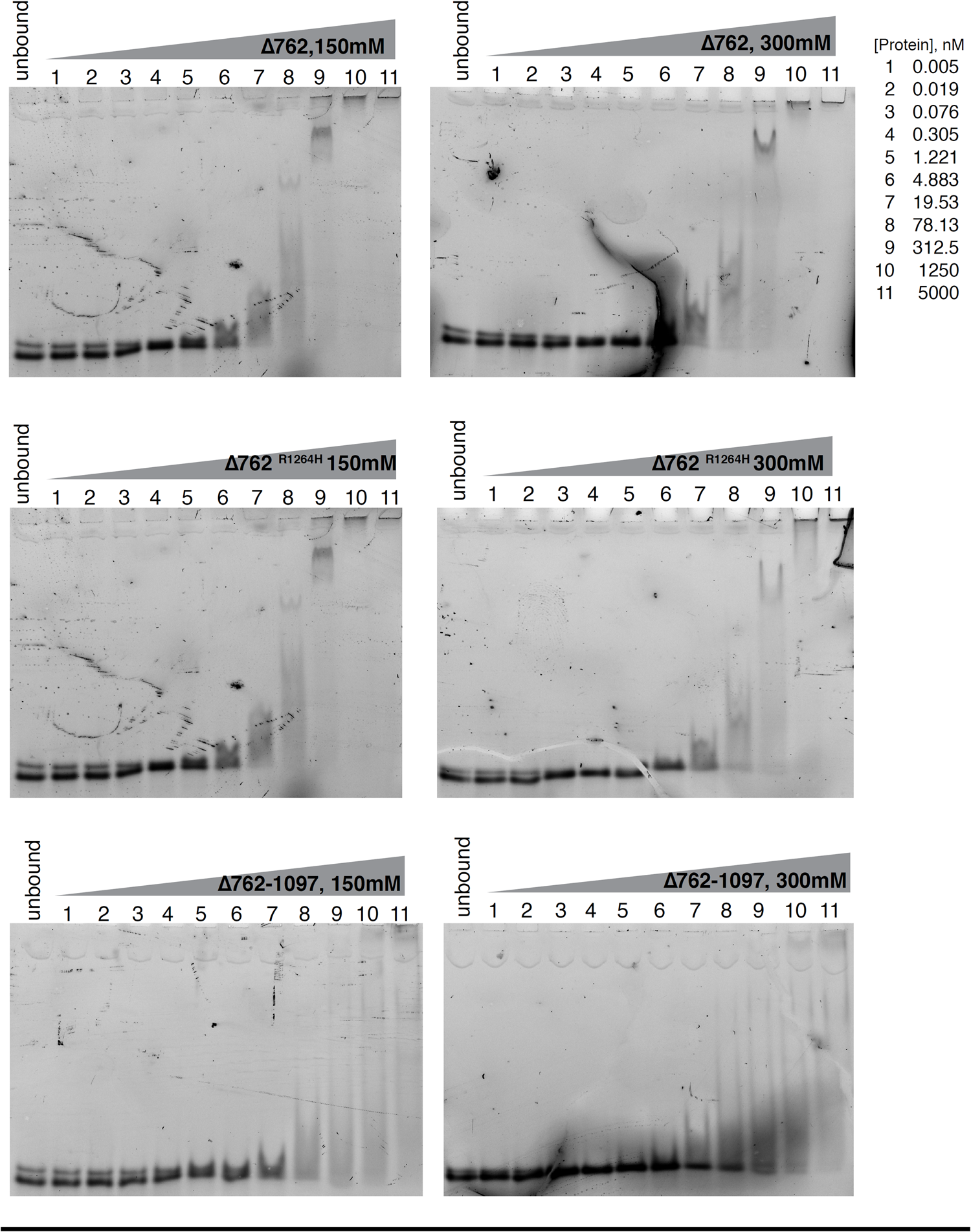
Representative native electrophoretic mobility shift assays (EMSA) with TERRA oligonucleotides at 2.5 nM and increasing amounts of the indicated recombinant proteins. Unbound indicates the free probe without protein added. EMSAs at 150 mM or 300 mM KCl are shown.

**Figure S5.**
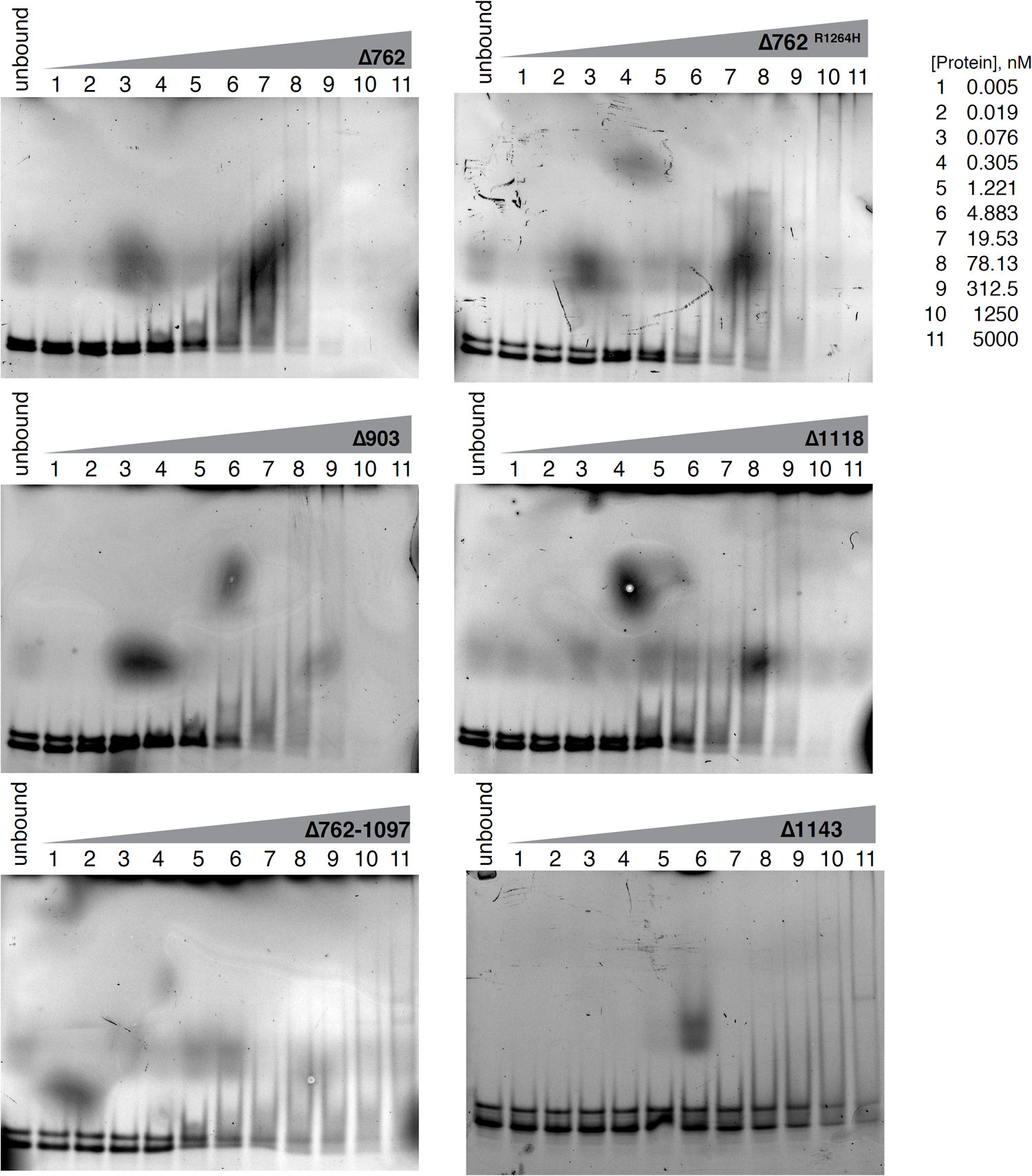
Representative native electrophoretic mobility shift assays (EMSA) with TERRA oligonucleotides at 2.5 nM and 150 mM KCl. Increasing amounts of the indicated recombinant RTEL1 truncated proteins were added. Unbound indicates the free probe without protein added. Negligible binding was observed for RTEL1^Δ1143^ at 5 *µ*M protein concentration.

**Figure S6.**
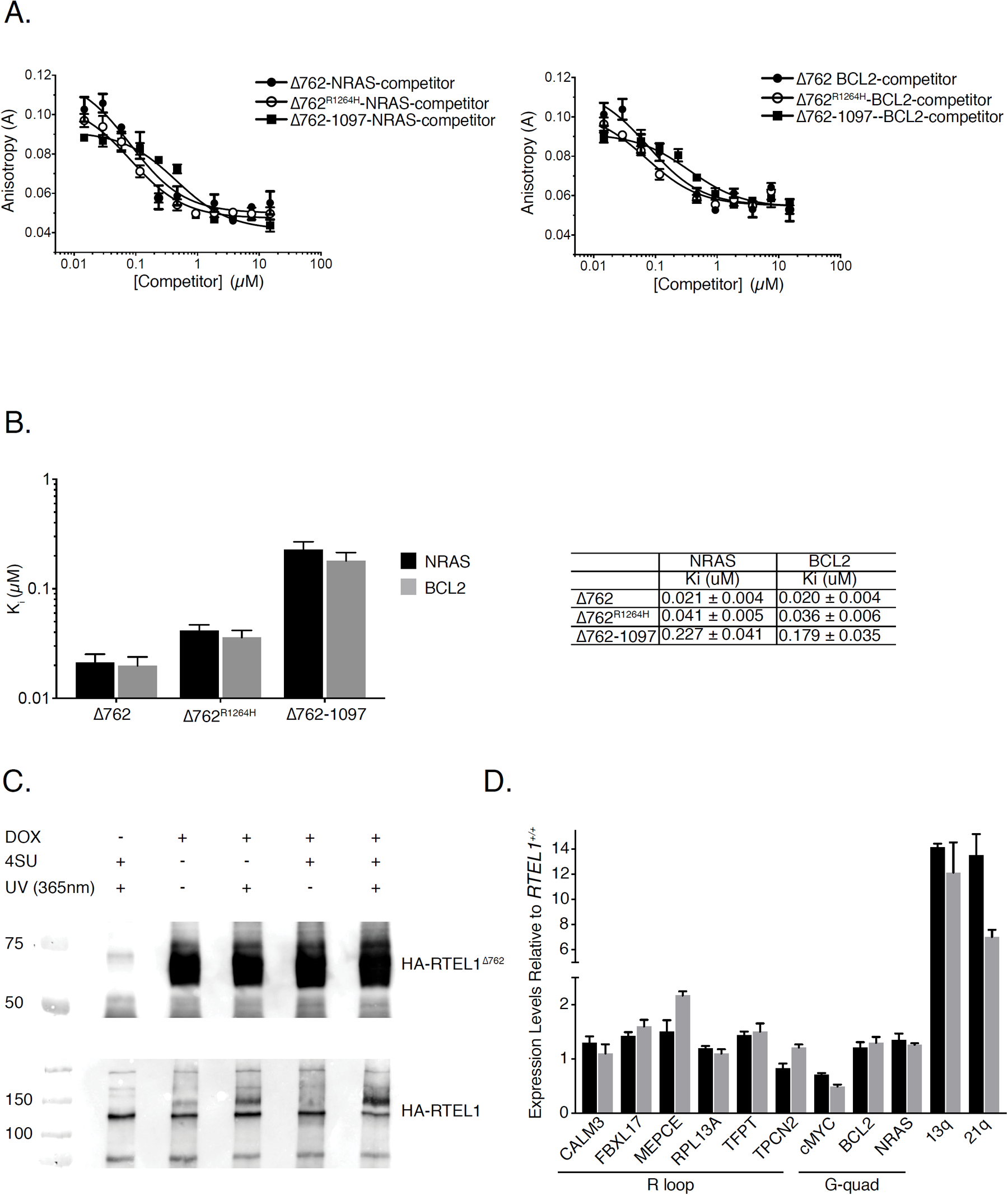
RTEL1 binding to NRAS and BCL2 oligonucleotides was monitored by fluorescence anisotropy. (A) Increasing concentrations of the indicated oligonucleotides were added to reactions containing the indicated RTEL1 proteins and a 24mer FAM-TERRA-MUT RNA. Binding curves used to calculate apparent dissociation constants (Ki) are shown. (B) Bar graphs show apparent dissociation constants (Ki) derived by competition experiments in A. All binding assays were conducted in triplicate and mean and standard deviation are shown. (C) Western blot analysis of cross-linked RNA-protein immunoprecipitates shown in Figure 2E. (D) Expression levels of non-telomere targets predicted to form R-loops and G-quadruplexes measured by qRT-PCR. No significant differences were observed in RTEL1-KO cells compared to wild type HEK293 cells in three independent experiments. Mean and standard deviation are shown.

**Figure S7.**
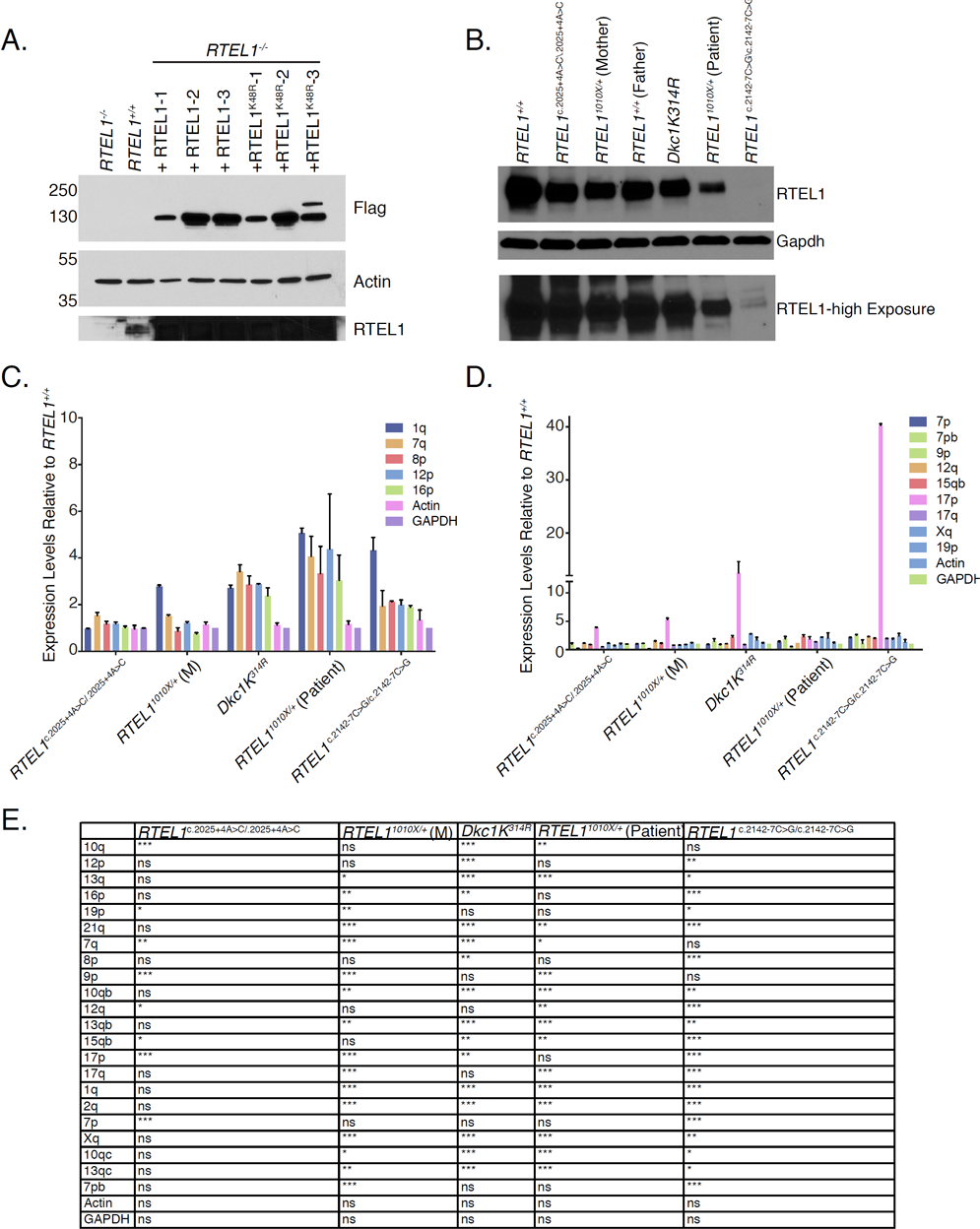
(A) Western blot of RTEL1 protein levels in wild type, RTEL1-KO, and three independent clones of *RTEL1-KO HEK293* cells reconstituted with FLAG-RTEL1 and FLAG-RTEL1^K48R^. Actin is shown as a loading control. (B) Endogenous RTEL1 protein levels were measured by western blotting in a panel of patient derived LCL cell lines. TERRA levels are elevated in cells derived from a panel of patient derived LCL lines. (C) Chromosome ends elevated 2.5-5 fold relative to wild type levels are shown. D) Chromosome ends elevated 2.5 fold or under relative to wild type levels are shown and levels for telomeric end 17p which is CTCF regulated is also shown (Beishline, Vladimirova et al. 2017). Levels were measured by qRT-PCRValues are sample averages of at least 3 experimental repeats. (E) Calculated p values for TERRA expression measured by qRT-PCR in three independent experiments shown including p values for Figure 3E (*p < 0.05, **p <0.01, ***p < 0.001, t-test).

**Figure S8.**
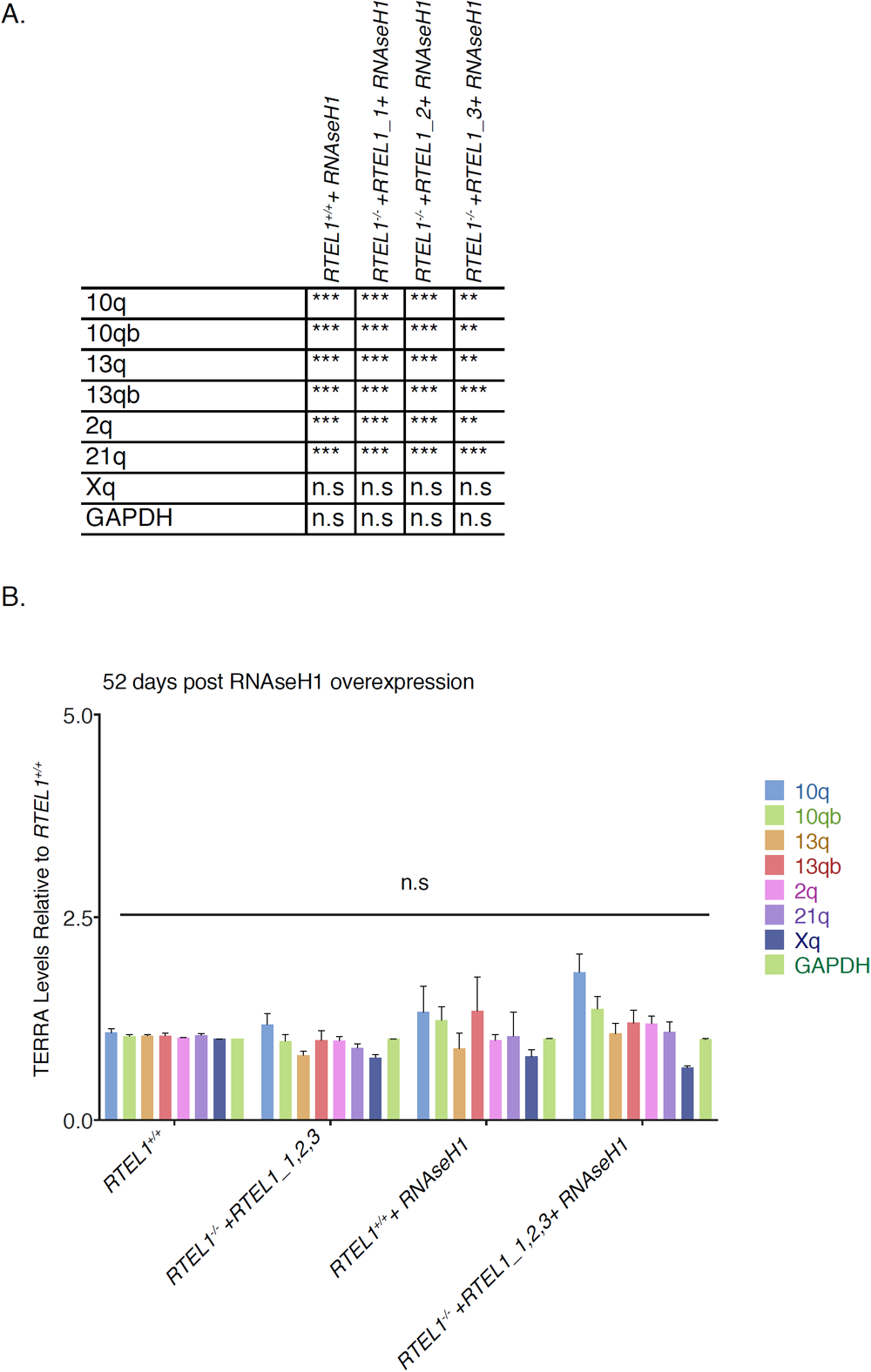
(A) Calculated p values for TERRA expression measured by qRT-PCR in three independent experiments shown in Figure 5B (*p < 0.05, **p <0.01, ***p < 0.001, t-test). (B) TERRA levels 52 days post *RNAse H1* overexpression. TERRA levels of indicated chromosome ends were tested in cells transfected with *RNAse H1-GFP* 52 days post G418 selection. TERRA levels were measured by qRT-PCR. Three independent cell lines complemented with RTEL1 were averaged for analysis. Values are sample averages of at least 3 experimental repeats (*p < 0.05, **p <0.01, ***p < 0.001, t-test).

